# Metabolic and transcriptional responses reveal co-option of symbiotic processes during early soybean-soybean cyst nematode interaction

**DOI:** 10.64898/2026.10.23.761348

**Authors:** Megha Rai, Esmaeil Miraeiz, Mainak Dutta, Vidhi Desai, Amit Rai, Matthew Hudson

**Author notes:** These authors contributed equally (Megha Rai, Esmaeil Miraeiz). Correspondence: Matthew Hudson [ ].

## Abstract

The biochemical mechanisms by which virulent populations of soybean cyst nematode (*Heterodera glycines*, SCN) overcome host resistance are critical to understanding this economically important pathogen. Here, we deployed metabolic and transcriptome profiling of soybean roots during the SCN infection and establishment process to investigate the molecular and biochemical mechanisms of pathogen establishment. Across samples from a time series spanning nematode invasion to syncytium maturation, we demonstrate that compatible SCN infection does not bypass host immunity but systematically remodels it. In a compatible interaction between soybean carrying the high-copy *Rhg1* resistance locus challenged with HG Type 2.5.7 SCN virulent on this host, we identify a 72-metabolite defense signature spanning isoflavonoids, triterpenoid saponins, and phenylpropanoids that persist across all infection stages. Despite activation of this metabolic defense signature by the pathogen on infection, defense transcription is near silence by 3 days post-infection, while defense metabolites persist, revealing a critical decoupling of transcriptional and metabolic defense programs at the precise window of syncytium initiation. Suppression of phenylpropanoid accumulation occurs downstream of PAL transcription, indicating a block at the metabolic level invisible to transcriptomics alone. By 5 days post-infection, cytokinin accumulation coincides with nodulin gene induction and phospholipid remodeling consistent with activation of the legume symbiosis program during syncytium maturation. Together, these findings establish that compatibility arises not from failed recognition but from progressive immune dismantling, including suppression of defense chemistry and co-option of the legume-rhizobium symbiosis machinery.

## INTRODUCTION

The soybean cyst nematode (*Heterodera glycines*; SCN) causes greater annual yield losses in soybean than any other single pathogen, with U.S. losses estimated to exceed $1.5 billion per year (Koenning and Wrather, 2010). SCN is an obligate biotroph that transforms host root cells into a syncytium, which is a multinucleate metabolic sink sustaining nematode growth and reproduction throughout the infection cycle (Gheysen and Mitchum, 2011; Jones and Northcote, 1972). The cellular remodeling required for syncytium formation shares mechanistic parallels with legume–rhizobium symbiosis: both involve cell wall dissolution, membrane proliferation, nodulin gene induction, and cytokinin-mediated developmental reprogramming (Favery et al., 2002; Redding et al., 2018). Consistent with this, early physiological and molecular work established that nematode infection activates nodulation associated gene expression in soybean roots (McGinnity et al., 1980; Redding *et al*., 2018), suggesting that parasitic nematodes may co-opt conserved symbiotic machinery for feeding site development. While transcriptional reprogramming is well-documented (Matthews et al., 2013), the coordination between these genetic changes and the resulting metabolic consequences remains a critical area for further investigation.

Resistance to SCN in U.S. soybean production has relied for decades on the *Rhg1* locus derived from PI 88788, which encodes *α-SNAP* and *NSF* proteins that disrupt syncytium function in incompatible interactions (Bayless et al., 2016; Cook et al., 2012; Kandoth et al., 2011). Virulent SCN populations capable of reproducing on high-copy *Rhg1* genotypes are now widespread across major soybean-producing states, undermining the most broadly deployed resistance source in the crop (McCarville et al., 2017; Miraeiz et al., 2026a; Miraeiz et al., 2026b; Mitchum, 2016). Whether virulent populations overcome *Rhg1*-mediated resistance by evading host recognition entirely, or by dismantling an already activated defense response at the metabolic and hormonal level, remains unresolved. This distinction has direct consequences for how durable resistance should be engineered.

Transcriptomic studies have established that early SCN infection triggers rapid defense gene induction followed by progressive transcriptional attenuation as the syncytium matures (Chaiprom et al., 2024; Ithal et al., 2007; Mitchum and Baum, 2008; Nissan et al., 2025; Zhang et al., 2026). Yet mRNA abundance cannot reveal whether these transcriptional changes translate into bioactive metabolite accumulation, which represents the actual chemical output that determines whether a host cell resists, dies, or is remodeled into a feeding site. This distinction is critical for SCN, where resistance is associated with rapid accumulation of isoflavonoid phytoalexins such as glyceollin, which are directly toxic to infective juveniles, while compatible interactions are characterized by disruption of defense chemistry at the metabolic level, independent of transcriptional changes (Huang and Barker, 1991; Neupane et al., 2019). Post-translational regulation, enzymatic competition, substrate channeling, and compartmentalization all shape metabolic output independently of transcription (Fernie and Morgan, 2013; Rai and Saito, 2016; Rai et al., 2017), and SCN effectors act at the protein level to redirect host cell fate (Mitchum et al., 2013). Phytohormone signaling further connects transcriptional responses to metabolic outputs. Salicylic acid contributes to resistance against sedentary endoparasites, jasmonate mediates systemic defense, and cytokinin promotes cell division and nodulin gene expression during symbiosis (Dehio and de Bruijn, 1992; Dong and Hudson, 2022; Sikder et al., 2021). Resolving whether compatibility arises from a failure of the host to recognize the pathogen, from a failure to activate a response at the transcriptional level, or from post-transcriptional dismantling of an already-activated defense therefore requires integrating metabolomics with transcriptomics across the infection timeline.

To resolve the metabolic and hormonal dimensions of SCN infection beyond what transcriptomics can reveal, we integrated metabolomics and transcriptomics at 1-, 3-, and 5-days post-infection (DPI) **(Figure 1)**, spanning nematode invasion, syncytium initiation, and feeding site maturation, in soybean carrying a high-copy *Rhg1* locus challenged with an SCN HG Type 2.5.7 population, a pathogen type which has overcome this type of resistance and become virulent on this host, creating an increasing problem for soybean agriculture in many areas (Chen and Potter, 2025). Our data reveals that this virulent SCN population does not evade host immune responses but remodels it into a compatible interaction. Defense metabolites are produced, but their biosynthesis is blocked post-transcriptionally at the phenylpropanoid entry point. Jasmonate defense is constitutively suppressed from the moment of infection, tryptophan is diverted away from auxin biosynthesis, and by 5 DPI, cytokinin accumulation coincides with nodulin induction, implicating activation of the legume-rhizobia symbiosis program during syncytium maturation. Together, these findings reframe compatibility as a progressive, multi-pathway dismantling of an activated defense, revealing that durable SCN resistance requires protecting the biochemical execution of immunity, not merely its initiation.

**Figure 1.**
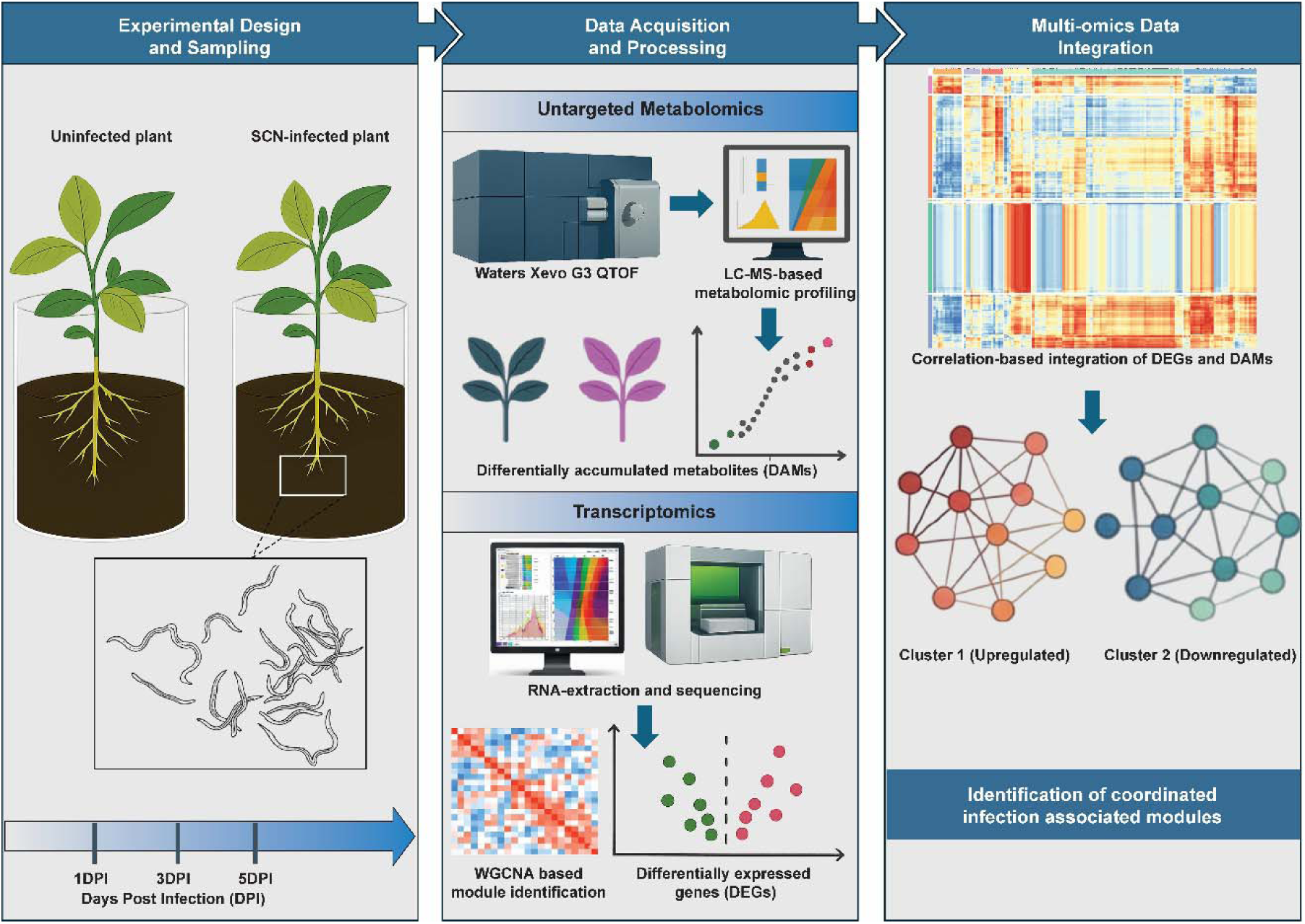
Experimental design and integrative multi-omics workflow for profiling early soybean responses to *Heterodera glycines* infection. Roots of Soybean roots were sampled at 1-, 3-, and 5-days post-inoculation (DPI) from both SCN-infected and non-infected plants to capture nematode penetration, syncytium initiation, and feeding site maturation. Untargeted metabolomics (Waters Xevo G3 QTof LC–MS) and strand-specific RNA sequencing were performed to identify differentially accumulated metabolites (DAMs) and differentially expressed genes (DEGs). Weighted gene co-expression network analysis (WGCNA) resolved transcriptional modules, while correlation-based integration of DEGs and DAMs identified coordinated infection-associated gene–metabolite clusters, including upregulated and downregulated programs.

## RESULTS

### Untargeted metabolomics reveals a stable 72-metabolite defense signature that persists throughout SCN infection

Soybean roots collected at 1-, 3-, and 5-days post-inoculation (DPI) *with Heterodera glycines*, together with non-infected controls, were subjected to untargeted LC-MS metabolomic profiling using Waters Xevo G3 Q-ToF mass spectrometer in positive ionization mode. Quality control and feature extraction using MS-DIAL (Tsugawa et al., 2015) yielded 34,695 mass features in positive ionization mode **(Supplementary Table 1)**, with high reproducibility across biological replicates confirmed by total ion chromatogram alignment **(Supplementary Figure S1)**. Unsupervised principal component analysis (PCA) revealed substantial overlap between infected and non-infected samples at all three time points, indicating that global metabolic variance was not dominated by infection status alone **(Supplementary Figure S2A)**. KEGG pathway mapping assigned chemical identities to 865 features **(Supplementary Table 2)**, with phenylpropanoid biosynthesis, isoflavonoid biosynthesis, and flavonoid biosynthesis among the top-represented pathways **(Supplementary Figure 2B)**, consistent with the central role of these pathways in legume defense against soil-borne pathogens (Huang and Barker, 1991; Neupane *et al*., 2019). Integrated annotation using MS-DIAL, MS-FINDER (Tsugawa et al., 2016), and SIRIUS (Ludwig et al., 2020) identified 196 metabolites across all samples **(Supplementary Table 3)**, including isoflavonoids, phenylpropanoids, triterpenoid saponins, iridoid glycosides, and lipid-associated metabolites.

Pan-timepoint OPLS-DA integrating all infected versus non-infected samples across the three time-point course discriminated infected from non-infected roots **(Figure 2A)**, identifying 816 differentially accumulated features (**Figure 2B**; **Supplementary Table 4**). Among the top-ranked discriminatory features were corylin, a prenylated isoflavonoid belonging to the class of soybean phytoalexins with well-documented roles in defense against soil-borne pathogens (Huang and Barker, 1991; Neupane *et al*., 2019) and acylated soysapogenol esters representing triterpenoid saponin derivatives with established role in soybean defense (Yates et al., 2021) **(Figure 2C)**. Of these 816 discriminatory features, 72 high-confidence plant-derived metabolites with confirmed structural annotation, selected based on annotation confidence and reproducibility and spanning six biochemical classes defined by their chemical ontology **(Figure 2D; Supplementary Table 5)**.

**Figure 2.**
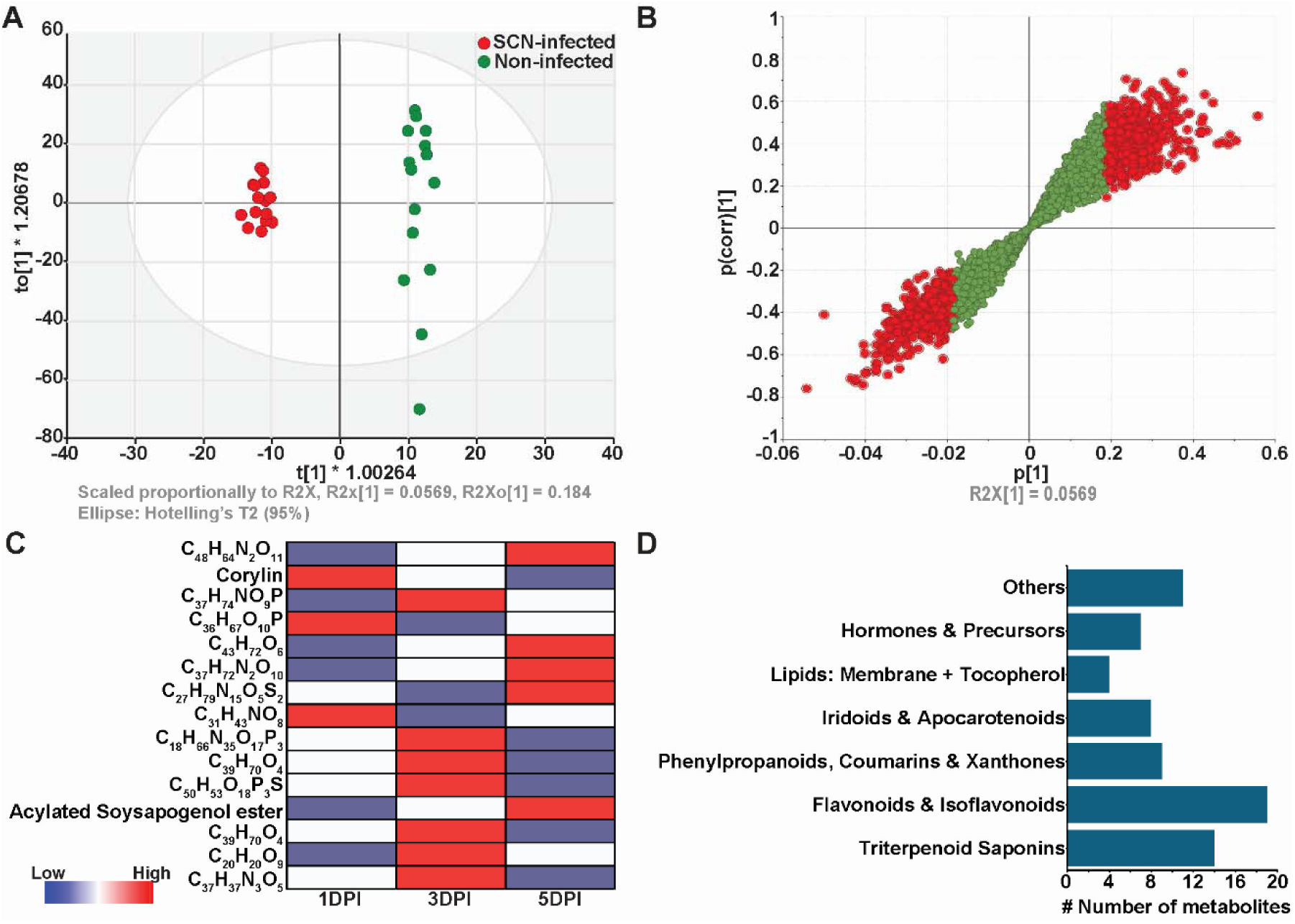
A stable, multi-class metabolic signature defines soybean’s systemic response to SCN infection. **(A)** Pan-timepoint O2PLS-DA score plot showing clear separation between all SCN-infected and non-infected root samples across 1–5 days post-inoculation (DPI), revealing a consistent infection-associated metabolic signature. **(B)** S-plot from the pan-timepoint O2PLS-DA model highlighting metabolites with high covariance (x-axis) and correlation (y-axis) driving the infected vs. non-infected separation. Metabolites highlighted in red represent 817 discriminatory features. **(C)** Heatmap of the top 15 metabolites contributing most to the infection signature (based on VIP > 1.5), showing absolute log₂ fold-changes (Infected/Non-infected) at 1, 3, and 5 DPI. Red = induced, blue = suppressed. **(D)** Biochemical classification of 65 high-confidence annotated metabolites among the 817 discriminatory features from the S-plot, selected based on structural confidence and relevance to plant–nematode interactions.

This core 72-metabolite signature included triterpenoid saponins including medicoside C, acetylsoyasaponin A6, yamogenin tetroside B, and oleanolic acid glycosides, which functions as membrane-disrupting antimicrobial compounds and have been reported at nematode infection site in legumes (Yates *et al*., 2021); iridoid glycosides including amarogentin, an iridoid glycoside and a member of the secoiridoid subclass of terpenoid secondary metabolites; and phenylpropanoid-derived metabolites including corylin, neobavaisoflavone, coumarin, kaempferol 3-α-L-arabinopyranoside, and luteolin 7-glucoside-4′-(Z-2-methyl-2-butenoate), all members of the isoflavonoid and flavonoid branches of phenylpropanoid metabolism that are well-established as primary soybean defenses against *H. glycines* (Huang and Barker, 1991; Neupane et al., 2019) **(Figure 2D; Supplementary Table 5)**. Lignans are phenylpropanoid-derived secondary metabolites that function primarily in plant defense, contributing to protection against pathogens, herbivores, and environmental stresses (Singh et al., 2024). Lirioresinol A, a syringaresinol-type lignan, was depleted at 1 DPI (0.25-fold relative to controls) and remained reduced at 3 DPI, before recovering to 1.7-fold above control levels by 5 DPI **(Supplementary Table 5)**. This biphasic pattern suggests that phenylpropanoid-derived metabolism is temporally modulated during infection, a pathway that also supplies precursors for lignin biosynthesis.

Hormone-associated metabolites showed coordinated changes across all infection stages: methyl dihydrojasmonate was suppressed 1.8–2.4-fold and abscisic acid aldehyde was progressively suppressed from 1.3– fold at 1 DPI to 3.3-fold at 5 DPI **(Supplementary Table 5)**. Methyl dihydrojasmonate is a derivative of the jasmonate signaling pathway, which mediates systemic defense responses against soil-borne parasites in soybean (Nahar et al., 2011; Zhang et al., 2017). Its consistent suppression across all three infection stages indicates that jasmonate-mediated defense is not maintained during compatible SCN infection. Mevalonic acid lactone, the committed product of the mevalonate pathway and precursor for terpenoid, cytokinin, and sterol biosynthesis, was elevated from 1.9-fold at 1 DPI to 1.5-fold at 5 DPI **(Supplementary Table 5)**, indicating sustained upregulation of isoprenoid biosynthesis throughout the infection cycle. Phosphatidylethanolamine (16:0/16:1) was consistently elevated from 1.7-fold at 1 DPI to 2.4-fold at 5 DPI **(Supplementary Table 5)**. Phosphatidylethanolamines are major structural components of plant membranes and accumulate during the membrane proliferation that characterizes both nematode syncytium development and the symbiotic interface in legume nodules (Catalano et al., 2007; Ivanov et al., 2012). The persistence of this 72-metabolite signature, including defense-associated isoflavonoids, antimicrobial saponins, and membrane-associated lipids, across all infection stages indicates that SCN establishes parasitism within an already biochemically active host environment rather than by suppressing constitutive chemical defenses.

**Table 1.** Overview of metabolomic feature detection and annotation depth.

| Description | Metrics |
| --- | --- |
| Total mass features detected | 34,695 |
| Mass Features with MS/MS | 6176 |
| Annotated metabolites | 307 |
| Metabolites mapped to KEGG pathway based on *m/z* | 865 |

### Temporal metabolomics identifies suppressed phenylpropanoid accumulation as a defining feature of compatible SCN infection

Time-point-resolved OPLS-DA identified 706, 779, and 667 differentially accumulated metabolites (DAMs) at 1-, 3-, and 5-DPI respectively **(Figure 3A; Supplementary Table 6)**. Overlapping set analysis demonstrated limited overlap among time points, with only 24 DAMs shared across all three stages (Fig. 3a), indicating that metabolic reprogramming is predominantly stage-specific. Of the 93 high-confidence annotated plant metabolites identified across stage-specific DAMs, 31 overlapped with the 72 core metabolites from the pan-timepoint analysis **(Supplementary Table 5)**. At 1 and 3 DPI, induced metabolites slightly outnumbered suppressed features, whereas at 5 DPI a marked shift toward widespread suppression was observed **(Figure 3B)**, consistent with a progressive transition from active chemical defense toward structural metabolic remodeling.

**Figure 3.**
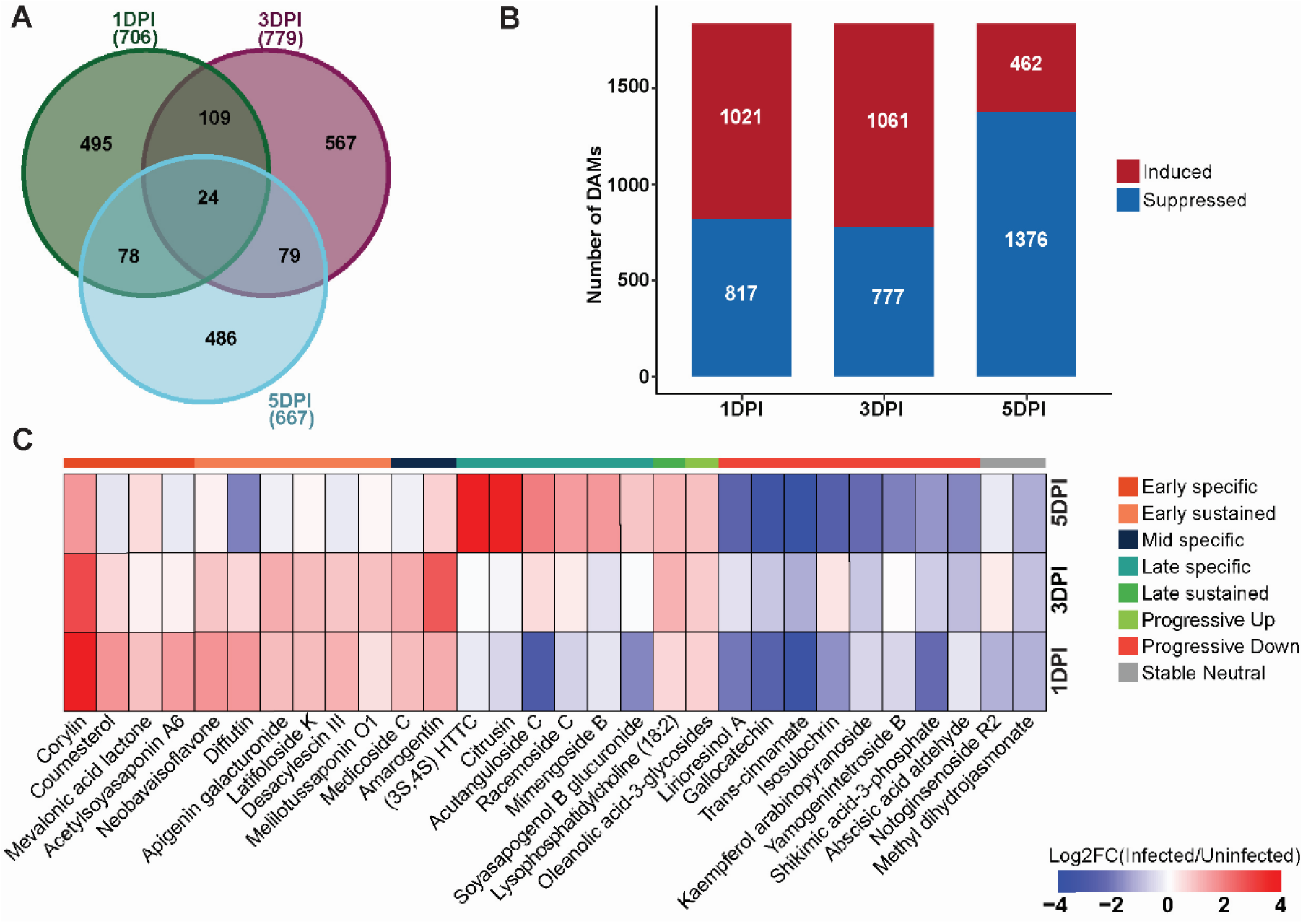
Temporal stratification of metabolic responses during early SCN infection. **(A)** Venn diagram showing overlap of differentially accumulated metabolites (DAMs) identified at 1-, 3-, and 5-days post-inoculation (DPI). **(B)** Number of induced and suppressed DAMs at each infection stage, classified based on the direction of log₂ fold change (infected vs non-infected). **(C)** Heatmap of 30 representative DAMs, selected based on magnitude of fold change (|log₂FC| ≥ 0.5) and biological relevance, illustrating distinct temporal accumulation patterns across infection stages. Colors indicate log₂ fold changes (infected/non-infected), with red denoting induction and blue denoting suppression.

Several metabolites showed persistent elevation across all three infection stages. Corylin, a pyranoisoflavonoid and member of the isoflavonoid phytoalexin pathway associated with defense against soil-borne pathogens (Huang and Barker, 1991; Neupane *et al*., 2019), was the most strongly and consistently induced metabolite, reaching 27.9-fold at 1 DPI and remaining elevated at 7.4-fold at 3 DPI and 3.0-fold at 5 DPI **(Figure 3C; Supplementary Table 5)**. Amarogentin, a secoiridoid glycoside, was elevated to 2.4-fold level at 1 DPI, peaked at 6.1-fold at 3 DPI, and remained 1.6-fold above controls at 5 DPI **(Fig. 3C; Supplementary Table 5)**. Secoiridoid glycosides have been implicated in plant defense through stress-induced accumulation, direct bioactivity against invading organisms, and interactions with root-associated microbial communities (Kubo et al., 1985; Tominaga et al., 2023). Thus, the accumulation of amarogentin at 3 DPI may reflect a defense-associated metabolic response to SCN invasion. Neobavaisoflavone, a prenylated isoflavonoid, was induced 3.1-fold at 1 DPI and remained slightly elevated through 3 and 5 DPI **(Figure 3C; Supplementary Table 5)**. Oleanolic acid 3-glycosides, a subclass of triterpenoid saponins, are structurally related to triterpenoid saponins reported to exhibit membrane-disrupting antimicrobial activity (Jiang et al., 2025), showed a progressive increase from 1.6-fold at 1 DPI to 1.9-fold at 5 DPI **(Supplementary Table 5)**. Lyso-phosphatidylcholine (18:2) was slightly elevated at 1 DPI (1.5-fold), increased to 2.3-fold at 3 DPI, and remained elevated at 2.1-fold at 5 DPI **(Fig. 3C; Supplementary Table 5)**. Lysophosphatidylcholines (LPCs) are bioactive lysophospholipids generated through phospholipase A₂ (PLA₂)-mediated hydrolysis of membrane phospholipids and are closely associated with phospholipid remodeling and membrane homeostasis (Kita et al., 2019; Murakami, 2023). Nematode-induced syncytium formation involves extensive cellular reprogramming, including profound changes in cytoplasmic organization, membrane-associated structures, and cellular transport capacity (Bohlmann and Sobczak, 2014; Siddique and Grundler, 2018). Thus, the accumulation of LPCs observed at nematode feeding sites may reflect altered membrane lipid metabolism and remodeling associated with syncytium development. Mevalonic acid lactone was elevated 1.9-fold at 1 DPI and remained elevated through 5 DPI **(Supplementary Table 5)**. As the committed product of the mevalonate pathway, its sustained elevation reflects upregulation of isoprenoid biosynthesis supplying precursors for cytokinin, sterol, and terpenoid synthesis, functional categories directly relevant to both the hormonal and structural metabolic remodeling observed at later infection stages.

A distinct group of metabolites showed strong early induction that declined progressively over the infection time course, consistent with a defense response that is activated but not maintained through the syncytium initiation window. Coumestrin was elevated to 3.5-fold at 1 DPI and 1.6-fold at 3 DPI before returning to control levels by 5 DPI **(Supplementary Table 5)**. Coumesterol, a coumestan phytoestrogen and member of the soybean isoflavonoid defense pathway induced during SCN infection (Chaiprom *et al*., 2024; Huang and Barker, 1991), peaked at 3.1-fold at 1 DPI and declined to 1.5-fold at 3 DPI and near-control levels by 5 DPI **(Fig. 3C; Supplementary Table 5)**. Diffutin was induced 3.0-fold at 1 DPI and declined to near-control levels by 5 DPI **(Fig. 3c; Supplementary Table 5)**. Medicoside C was elevated 2.1-fold at 1 DPI and 1.8-fold at 3 DPI before returning to control levels at 5 DPI **(Supplementary Table 5)**. Desacylescin III, a triterpene saponin, followed a similar trajectory, induced 2.3-fold at 1 DPI and 1.8-fold at 3 DPI before declining at 5 DPI **(Supplementary Table 5)**. Indolelactic acid, a shunt metabolite of the tryptophan-to-auxin biosynthetic pathway, was elevated 2.5-fold at 1 DPI and 1.6-fold at 3 DPI before returning to near-control levels by 5 DPI **(Supplementary Table 5; Supplementary Figure S5)**. Elevated indolelactic acid in the context of declining free IAA is consistent with enzymatic redirection of tryptophan metabolism away from productive auxin biosynthesis (Chopra et al., 2021), a post-transcriptional mechanism invisible to transcriptomics alone.

Phenylpropanoid pathway intermediates showed a distinctive pattern of progressive suppression that reveals a mechanistically important post-transcription block on defense biosynthesis. Trans-cinnamate, the direct product of phenylalanine ammonia-lyase (PAL) and the committed entry point of phenylpropanoid biosynthesis, was suppressed to 8% of non-infected control levels at 1 DPI, partially recovered to 43% at 3 DPI, and declined further to 6% of controls by 5 DPI **(Figure 3C; Supplementary Table 3)**. Shikimic acid 3-phosphate, an upstream intermediate of the shikimate pathway that feeds into phenylpropanoid and aromatic amino acid biosynthesis, followed a parallel trajectory, suppressed to 20% control levels at 1 DPI, partially recovering to 58% at 3 DPI, and declining again to 34% at 5 DPI **(Supplementary Table 3)**. Critically, PAL transcript levels showed no differential expression at any infection stage **(Supplementary Table 7)**, indicating that this suppression of phenylpropanoid branch operates post-transcriptionally. Methyl dihydrojasmonate was consistently depleted across all time points, at 45%, 54%, and 42% of non-infected control levels at 1, 3, and 5 DPI respectively **(Figure 3C; Supplementary Figure S6; Supplementary Table 5)**, indicating that jasmonate defense is constitutively attenuated from the moment of infection rather than progressively declining in response to syncytium establishment. Abscisic acid aldehyde was progressively suppressed from 1.3-fold at 1 DPI to 3.3-fold at 5 DPI **(Supplementary Table 5)**, consistent with broad suppression of hormone-mediated defense signaling across the infection time course.

At 5 DPI, the metabolic landscape was dominated by the emergence of structural and membrane-associated metabolites, reflecting the transition from active defense to feeding site maintenance. The oxylipin derivative (3S,4S)-3-hydroxytetradecane-1,3,4-tricarboxylic acid was near-control levels at 1 and 3 DPI but was strongly induced 16.8-fold at 5 DPI **(Supplementary Table 5)**. Oxylipins are lipid-derived signaling metabolites generated through lipoxygenase-mediated oxidation of polyunsaturated fatty acids and have roles in defense signaling and membrane remodeling in plants (Mosblech et al., 2009). Citrusin was suppressed at 1 DPI and near-control at 3 DPI before rising to 12.9-fold above controls at 5 DPI **(Supplementary Table 5)**. Acutanguloside C was suppressed at 1 DPI (13% of controls), returned to near-control levels at 3 DPI, and was induced 3.8-fold by 5 DPI **(Supplementary Table 5)**. Racemoside C rose to 2.8-fold above controls at 5 DPI **(Supplementary Table 5)**. Lirioresinol A recovered from suppression at 1 DPI to 1.7-fold above controls by 5 DPI **(Supplementary Data 5)**. The coordinated emergence of oxylipin derivatives and structural metabolites at 5 DPI, coinciding with the suppression of early defense isoflavonoids, reflects a metabolic transition consistent with the establishment of the mature syncytium as a sustained metabolic sink.

### RNA-seq confirms biphasic defense silencing and identifies nodulin induction coinciding with cytokinin accumulation at syncytium maturation

Strand-specific RNA-seq of 24 libraries, four biological replicates per condition per time point, generated 28.4–37.1 million paired-end reads per sample, of which >99.5% passed quality filtering **(Supplementary Figure S3A, B)**. GC content of cleaned libraries ranged from 44–45% **(Supplementary Figure S3C)**. Fragment length distributions were consistent across all samples **(Supplementary Figure S3D)**. A total of 46,253 genes were identified across all samples **(Supplementary Table 7)**. t-SNE analysis of the global transcriptome revealed clear separation between infected and non-infected samples across all time points **(Figure 4A)**. Differential expression analysis (|log₂FC| > 1, FDR < 0.05) identified 290 unique DEGs across the three time points, with the highest number at 1 DPI, a marked reduction at 3 DPI, and a partial resurgence at 5 DPI **(Figure 4B; Supplementary Table 8)**. The number of significantly enriched Gene Ontology (GO) terms followed the same pattern, being highest at 1 DPI, sharply reduced at 3 DPI, and increased again at 5 DPI **(Figure 4C; Supplementary Table 8)**.

**Figure 4.**
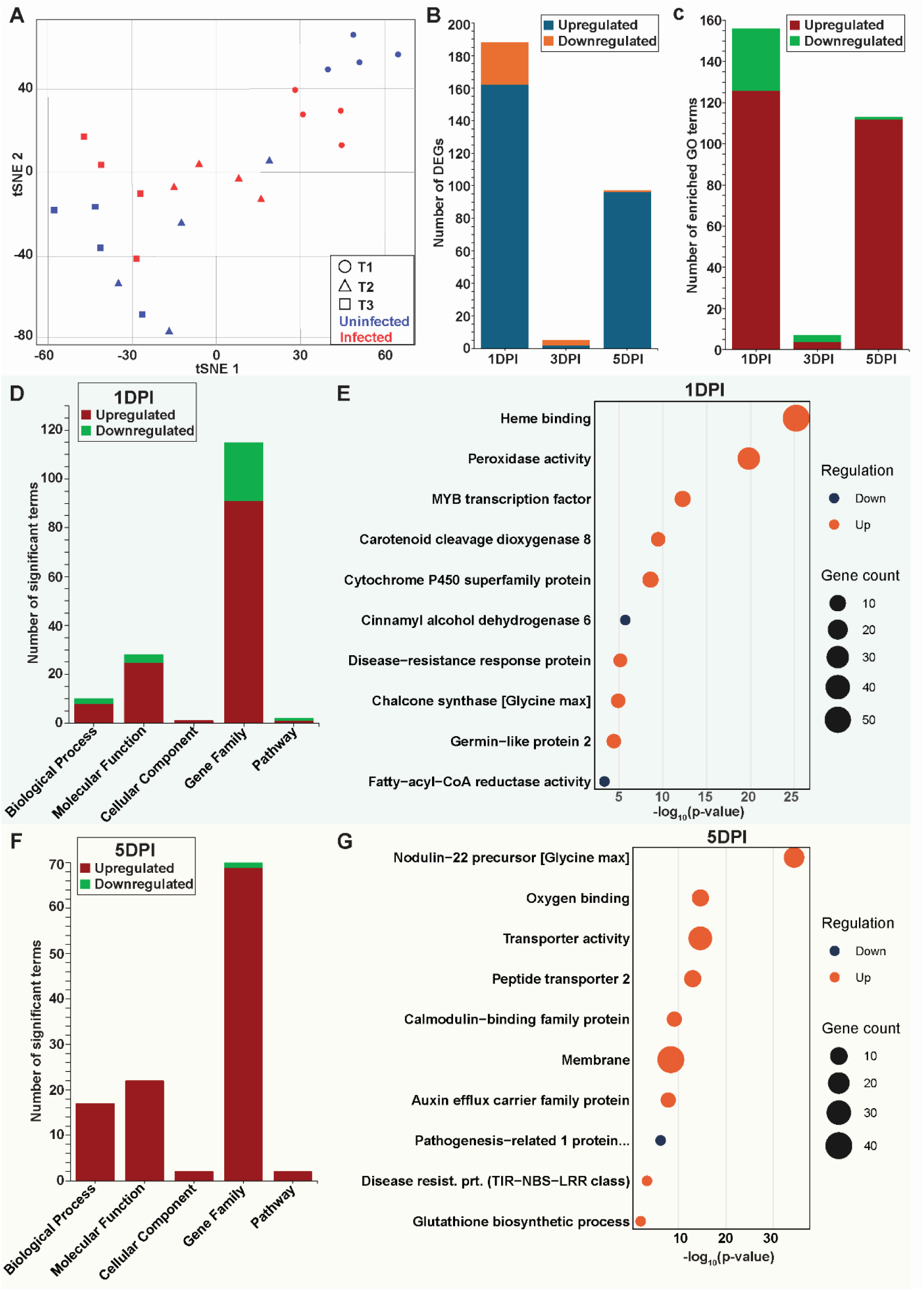
Stage-specific transcriptional reprogramming during early SCN infection. **(A)** t-SNE ordination of soybean root transcriptomes showing separation between SCN-infected and non-infected samples across 1-, 3-, and 5-days post-inoculation (DPI). **(B)** Number of differentially expressed genes (DEGs; |log₂FC| > 1, FDR < 0.05) identified at each time point, stratified by direction of regulation. **(C)** Number of significantly enriched Gene Ontology (GO) terms (P < 0.05) associated with up– and downregulated DEGs at each time point. **(D, F)** Distribution of enriched GO terms at 1 DPI and 5 DPI, respectively, grouped by ontology category (Biological Process, Molecular Function, Cellular Component, Gene Family, Pathway). **(E, G)** Top enriched GO terms and representative gene families at 1 DPI and 5 DPI, respectively, selected based on statistical significance (P < 0.05) and relevance to plant–nematode interactions.

At 1 DPI, upregulated DEGs were enriched for peroxidase activity, cytochrome P450 superfamily proteins, chalcone synthase, and carotenoid cleavage dioxygenase 8 **(Figure 4D, E; Supplementary Table 8)**. Chalcone synthases catalyze the first committed step of flavonoid and isoflavonoid biosynthesis (Anguraj Vadivel et al., 2021; Bian et al., 2018; Liu et al., 2025), and cytochrome P450 enzymes mediate downstream isoflavonoid hydroxylation (Wang et al., 2026). The coordinated induction of chalcone synthase and cytochrome P450s at 1 DPI is consistent with the strong corylin, coumesterol, and neobavaisoflavone accumulation identified in the metabolomics data, confirming that the early defense transcriptional response is functionally coupled to phenylpropanoid and isoflavonoid biosynthesis. Cinnamyl alcohol dehydrogenase 6 was simultaneously downregulated **(Supplementary Table 8)**, indicating that phenylpropanoid branch was directed toward soluble defense compounds rather than structural lignin polymerization at this stage.

By 3 DPI, defense transcription had collapsed to near-baseline: germin-like protein 2 was the sole significantly induced transcript (Supplementary Data 8). Germin-like proteins are apoplastic enzymes with superoxide dismutase or oxalate oxidase activity implicated in reactive oxygen species management during pathogen challenge (Dunwell et al., 2008). This near-complete transcriptional silencing at 3 DPI occurs precisely when the metabolomics data show that defense metabolites produced during the 1 DPI response persist without renewal, thus demonstrating a temporal decoupling of transcriptional activity from metabolite accumulation.

At 5 DPI, the transcriptional program shifted to nodulin-22 precursor induction, peptide transporter activity, membrane-associated functions, and oxygen-binding functions **(Figure 4F, G; Supplementary Table 8)**, while PR-1 was significantly downregulated **(Supplementary Figure 8)**. Nodulin proteins were originally characterized as legume-specific factors that establish and maintain the peribacteroid membrane during rhizobium symbiosis (Verma and Hong, 1996), and subsequent work established their expression at nematode feeding sites in both *Medicago truncatula* and soybean (Redding *et al*., 2018). The temporal specificity of nodulin-22 induction, which is absent at 1 DPI during invasion and present specifically at 5 DPI during active syncytium maturation, identifies symbiosis-associated transcription as a stage-specific feature of feeding site development rather than a non-specific response to cellular reorganization. Peptide transporters play roles in nitrogen assimilation and signaling across the symbiotic interface in legume nodules (Okumoto and Pilot, 2011), and their co-induction with nodulin-22 at 5 DPI indicates that multiple components of the legume symbiosis machinery are coordinately engaged during syncytium maturation. Notably, this transcriptional shift toward symbiosis-associated gene expression at 5 DPI coincides with the accumulation of lyso-phosphatidylcholine (18:2) and phosphatidylethanolamines identified in the metabolomics data. These membrane lipid species characterize the symbiotic interface in legume nodules (Catalano *et al*., 2007; Ivanov *et al*., 2012), thereby providing convergent metabolomic and transcriptomic evidence for symbiotic-type membrane reorganization during SCN syncytium maturation.

### Co-expression network analysis defines the transcriptional architecture of immune remodeling during SCN infection

WGCNA^31^ performed on 46,253 transcripts resolved 21 co-expression modules **(Figure 5A; Supplementary Table 9)**. Six modules, ME1, ME3, ME4, ME6, ME10, and ME16, exhibited clear differences in eigengene values between infected and non-infected samples across one or more time points **(Figure 5B)**. Of the 290 DEGs identified in the differential expression analysis, 252 (87%) were assigned to these six modules **(Supplementary Figure 9)**. ME1 and ME10 displayed elevated eigengene values specifically at 1 DPI in infected roots and were enriched for redox regulation, hormone biosynthesis, and primary metabolism **(Figure 5B, C; Supplementary Table 10)**, pathways consistent with the early defense transcriptional activation and isoflavonoid biosynthesis identified at 1 DPI. ME4 showed a pronounced increase in infected samples at 5 DPI and was enriched for membrane-associated transport and stress functions **(Figure 5B, C; Supplementary Data 10)**, consistent with the metabolic transition toward membrane remodeling and structural support observed at this stage. ME3 and ME6 exhibited higher eigengene values at 5 DPI in infected roots, with ME6 showing moderate elevation at 3 DPI as well **(Figure 5B)**. ME16 showed modest but consistent elevation at both 1 and 5 DPI while remaining near baseline at 3 DPI **(Figure 5B)**, suggesting a constitutively deployed infection-associated module whose activity at 3 DPI is not reflected in the DEG analysis.

**Figure 5.**
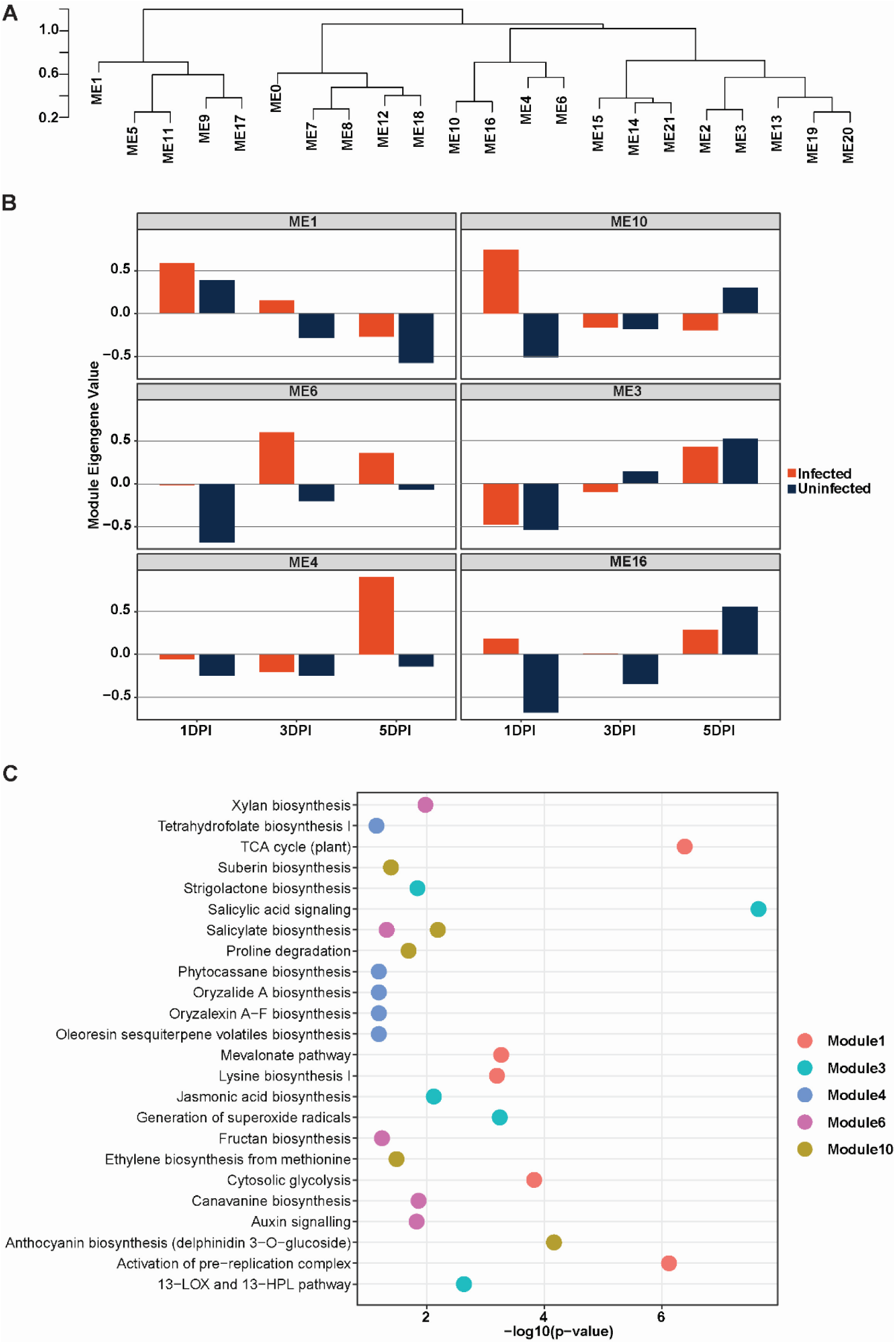
Weighted gene co-expression network analysis reveals temporally structured transcriptional responses during SCN infection. **(A)** Hierarchical clustering dendrogram of expressed genes showing module assignments (ME1–ME20). **(B)** Module eigengene values for six infection-responsive modules (ME1, ME10, ME6, ME3, ME4, and ME16). Bars represent mean eigengene values for infected (red) and uninfected (dark blue) samples at 1-, 3-, and 5-days post-inoculation (DPI). **(C)** Pathway enrichment analysis of the infection-responsive modules. Enriched pathways and biological processes are shown along the y-axis, with significance expressed as −log10(p-value). Colored points indicate the module in which each pathway was enriched.

### Gene–metabolite correlation analysis identifies coordinated transcriptional–metabolic modules during SCN infection

While transcriptomic and metabolomic analyses independently revealed strong differential regulation of transcription and metabolism during the time series, the extent to which transcriptional profiles are coordinated with metabolite accumulation remained unclear. To address this, correlation-based integration was performed using differentially expressed genes and differentially accumulated metabolites across infection stages **(Supplementary Table 11)**, enabling assessment of coordinated gene–metabolite behavior over time. Pearson correlation analysis between DEGs and DAMs across all infection stages resolved four gene clusters (GC1–GC4) and six metabolite clusters (MC1–MC6), forming a structured correlation network characterized by both positive and negative associations **(Supplementary Figure S4; Supplementary Table 12)**.

At 1 DPI, gene clusters GC2 and GC3 exhibited strong positive correlations with MC4 flavonol glycosides and MC5 isoflavonoid phytoalexins, and strong negative correlations with MC6 phospholipid-associated metabolites **(Figure 6; Supplementary Table 13)**. GC2 contains chalcone synthases, peroxidases, and WRKY transcription factors; GC3 contains MYB transcription factors, LRR receptor-like kinases, and cytochrome P450 enzymes. The positive correlation between these defense-associated gene clusters and the isoflavonoid and flavonol metabolite clusters at 1 DPI confirms that early defense gene activity is functionally coupled to the production of phenylpropanoid and isoflavonoid defense metabolites (Anguraj Vadivel *et al*., 2021; Bian *et al*., 2018; Liu *et al*., 2025; Wang *et al*., 2026), with chalcone synthase 5 showing a high-confidence hub correlation with corylin and polyphenol oxidase A1 with astilbin **(Supplementary Table 13)**.

**Figure 6.**
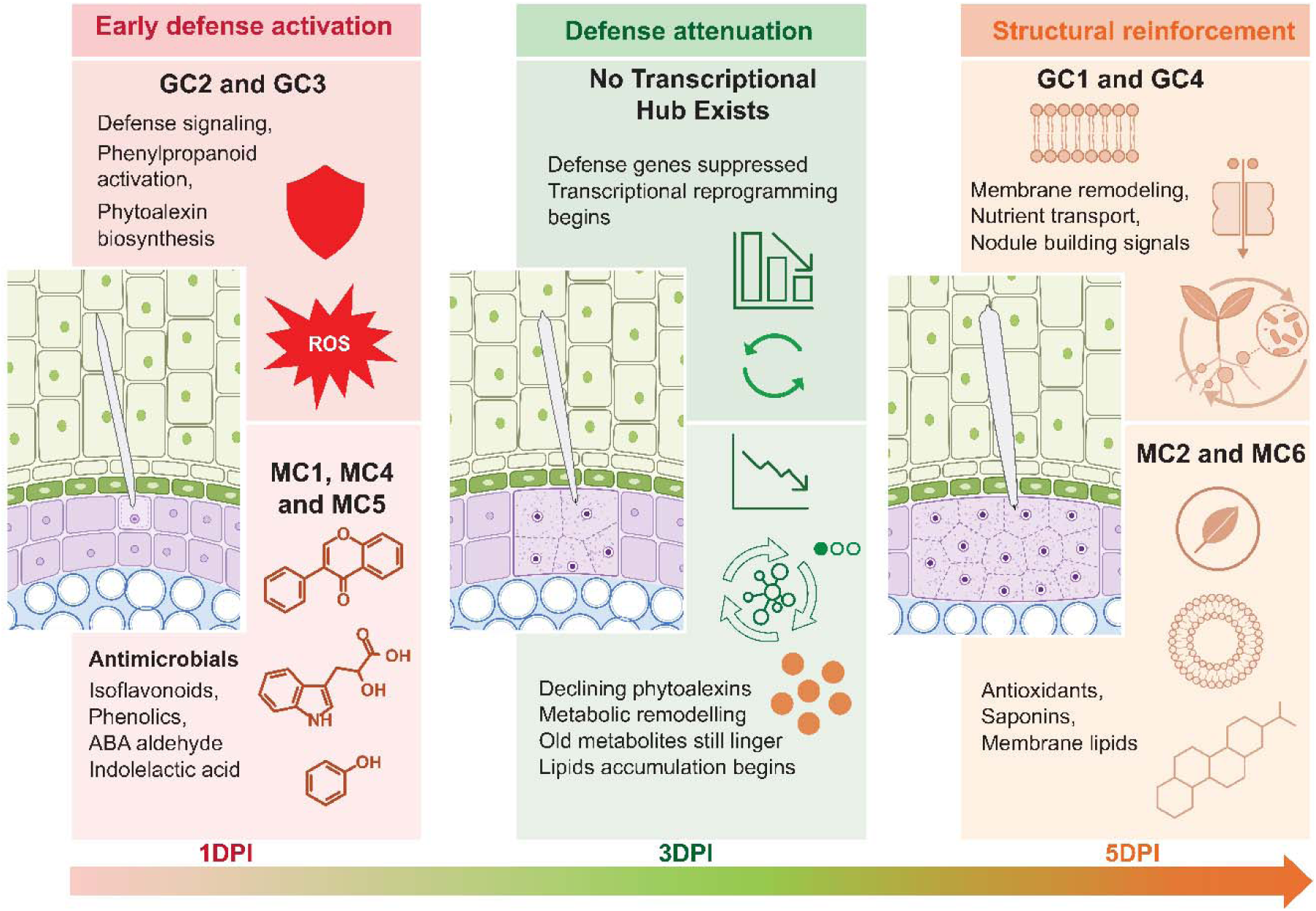
*Rhg1*-mediated partial resistance is characterized by phased transcriptional and metabolic reprogramming during soybean cyst nematode infection. Conceptual model integrating infection-induced transcriptomic changes (infected vs. non-infected roots) with significant gene–metabolite correlations (|r| ≥ 0.75, p < 0.05) across early (1 DPI), intermediate (3 DPI), and late (5 DPI) stages of infection. At 1 DPI, defense-associated gene modules show strong induction and connectivity with downstream metabolic clusters. At 3 DPI, transcriptional activity is reduced, while metabolite accumulation persists, consistent with a temporal lag between gene expression and metabolic output. At 5 DPI, coordinated accumulation of membrane-associated and nutrient reallocation metabolites predominates, indicating a transition toward cellular stabilization and sustained tolerance. Only infection-responsive genes and metabolites

At 3 DPI, correlations between GC2 and GC3 and their associated metabolite clusters were markedly reduced despite sustained accumulation of metabolites within MC4 and MC5 **(Figure 6, Supplementary Table 13; Supplementary Figure S6)**. This decoupling of gene cluster activity from metabolite accumulation at 3 DPI, which represents the same time-point at which transcription collapses to near-baseline, provides integrated transcriptome-metabolome evidence that defense chemistry at 3 DPI persists as a biochemical legacy of 1 DPI activity rather than through active transcriptional maintenance. The absence of transcriptional support for ongoing metabolite accumulation at 3 DPI is consistent with the post-transcriptional suppression of phenylpropanoid branch identified in the metabolomics data, in which PAL transcription is unaltered while trans-cinnamate output remains severely suppressed.

At 5 DPI, GC1, including genes like nodulin-22 precursor, peptide transporter, 2-isopropylmalate synthase, and non-symbiotic hemoglobin transcripts, enriched specifically at syncytium maturation, displayed strong positive correlations with MC2, a membrane-associated metabolite cluster dominated by lyso-phosphatidylcholine (18:2) and multiple phosphatidylethanolamine species **(Figure 6; Supplementary Table 13)**. Phosphatidylethanolamines and lysophosphatidylcholines are major components of the peribacteroid membrane that encapsulates nitrogen-fixing bacteroids in legume nodules (Catalano *et al*., 2007; Ivanov *et al*., 2012), and their accumulation has been specifically associated with the membrane proliferation events that accommodate symbiont expansion during nodule development. The positive correlation between GC1, a gene cluster whose transcriptional content directly reflects the legume symbiosis program, and MC2, a metabolite cluster reflecting membrane lipid remodeling of the type associated with symbiotic interface formation, provides integrated transcriptomic and metabolomic evidence that the symbiosis-associated developmental program engaged at 5 DPI is accompanied by corresponding changes in membrane lipid composition. GC4 showed positive correlations with MC6 structural metabolites and negative correlations with MC5 phytoalexins at 5 DPI **(Figure 6; Supplementary Table 13)**, consistent with the active suppression of phytoalexin biosynthesis as syncytium development proceeds. A progressive shift in expression of WRKY, MYB, and bHLH transcription factors, which were strongly induced at 1 DPI and largely suppressed by 5 DPI **(Supplementary Figure S6; Supplementary Table 13)**, mirrors the decline in phytoalexin accumulation and identifies these regulators as the most proximate transcriptional correlates of late-stage defense suppression. Gene–metabolite correlation connectivity across all clusters is summarized in Table 2.

**Table 2.** Gene–metabolite cluster connectivity exceeding 100 significant associations.

| Gene cluster | Metabolite cluster | Number of significant correlations |
| --- | --- | --- |
| 3 | 1 | 815 |
| 1 | 2 | 781 |
| 3 | 5 | 508 |
| 2 | 6 | 470 |
| 2 | 5 | 461 |
| 1 | 1 | 316 |
| 3 | 6 | 292 |
| 2 | 1 | 240 |
| 4 | 5 | 136 |
| 2 | 4 | 129 |
| 2 | 2 | 119 |
| 4 | 1 | 107 |
| 1 | 4 | 106 |
| 1 | 5 | 104 |
| 4 | 6 | 101 |

## DISCUSSION

This study establishes that compatible SCN infection in soybean carrying the high-copy *Rhg1* locus does not result from a failure of immune recognition. The host recognizes the nematode, mounts a substantial and chemically genuine defense response, and produces defense metabolites across all infection stages. Compatibility instead reflects the systematic dismantling of that defense through hormonal reprogramming, metabolic suppression, and biochemical remodeling that transcriptomics alone cannot reveal. Central among these is the engagement of the legume symbiosis program: by 5 DPI, cytokinin accumulation coincides with nodulin gene induction and phospholipid remodeling, revealing that SCN redirects the host’s own symbiotic developmental machinery toward feeding site construction while simultaneously dismantling immune outputs. This conclusion reframes how the field should understand *Rhg1* durability failure and defines mechanistic targets for resistance engineering that go beyond what single-locus strategies can provide.

The connection between sedentary endoparasitic nematodes and the legume symbiosis program has deep roots in the physiological literature that has not being fully understood. Early work established that SCN infection elicits nodulation-associated physiological responses in soybean roots (Huang et al., 1984; Huang and Barker, 1983), and that the cellular reorganization required for syncytium formation bears striking parallels to the membrane proliferation and cytoplasmic reorganization that define the symbiotic interface in legume nodules (Koltai et al., 2001). Subsequent molecular work by Favery et al. demonstrated that *ENOD40* and *CCS52a*, genes defining the endosymbiotic program, are expressed at nematode feeding sites in *Medicago truncatula* (Favery et al., 2002), and Redding et al. documented multiple nodulation gene upregulations during reniform nematode feeding site establishment in soybean specifically (Redding *et al*., 2018). Our integrated data advance this picture in two critical ways: they establish the precise temporal window of symbiosis-associated transcription as active syncytium maturation at 5 DPI rather than initial invasion, and they provide the first metabolomic evidence that this transcriptional shift is accompanied by membrane lipid remodeling of the type associated with symbiotic interface formation, as demonstrated by the positive correlation between GC1 and MC2 in the gene-metabolite cluster analysis.

The progressive collapse of trans-cinnamate, 8% of controls at 1 DPI, partial recovery at 3 DPI, then 6% of controls by 5 DPI, without any differential change in PAL transcript abundance is the most mechanistically significant metabolomic finding of this study. It demonstrates that SCN disrupts phenylpropanoid branch post-transcriptionally, at the first committed step of the pathway, while leaving the transcriptional machinery intact. The biosynthetic genes are expressed but the metabolic output is suppressed. This mechanism explains why transcriptomic approaches have systematically underestimated SCN’s manipulation of host defense: the pathway appears active at the mRNA level while its chemical output is silenced at the metabolite level. This finding resolves a long-standing paradox in SCN-soybean literature. Huang and Barker established that susceptible interactions are characterized by reduced glyceollin accumulation despite apparent upstream gene activity, attributing this to post-transcriptional regulation (Huang and Barker, 1991), a hypothesis our data now support at molecular resolution. The parallel suppression of shikimic acid 3-phosphate across all infection stages further indicates that the block operates across multiple entry points of the phenylpropanoid pathway rather than at a single enzymatic step. Critically, this suppression is progressively reinforced as the syncytium matures, becoming most severe at 5 DPI, which is consistent with ongoing effector activity that scales with feeding site establishment.

The parallel disruption of auxin homeostasis reinforces this picture of coordinated post-transcriptional immune remodeling. Although tryptophan and free IAA did not meet the threshold for differential accumulation in the OPLS-DA analysis, indolelactic acid, a confirmed differentially accumulated metabolite and a competitive shunt of the tryptophan-to-IAA biosynthetic route, was elevated 2.5-fold at 1 DPI and remained elevated at 3 DPI before returning to near-control levels by 5 DPI **(Supplementary Figure S5)**. The pattern is consistent with enzymatic redirection of tryptophan away from productive IAA synthesis toward the indolelactic acid shunt, effectively decoupling precursor availability from auxin output. This is further supported by the finding that cyst nematodes redirect host indole metabolism via NADPH oxidase-mediated ROS to promote infection (Chopra *et al*., 2021). SCN effectors are known to modulate auxin signaling to reprogram vascular parenchyma cells into the feeding site (Grunewald et al., 2009; Mitchum *et al*., 2013), and the indolelactic acid accumulation observed here, a post-transcriptional metabolic event invisible to transcriptomics, is consistent with active manipulation of auxin homeostasis to maintain a parasitism-permissive cellular environment. Whether this diversion reflects nematode-driven suppression of auxin biosynthesis or a host counter-strategy to limit auxin-dependent cell reprogramming cannot be resolved from the current data, but the metabolic outcome constitutes a precise disruption of the auxin signaling axis that transcriptomics would not detect.

At 3 DPI, transcriptional activity collapses to near-silence, with only germin-like protein 2 and a ribosomal protein among the significantly induced transcripts. Yet the isoflavonoids, saponins, and phenylpropanoids accumulated at 1 DPI remain elevated, creating a state in which defense chemistry persists without transcriptional renewal, thus defining the critical window of compatibility establishment. This 3 DPI configuration, active residual metabolites, silenced transcription, decoupled gene-metabolite correlations, is the molecular tipping point between resistance and established parasitism. In functionally resistant interactions, plants maintain coordinated transcriptional and metabolic defense responses through the early stages of nematode attack (Chaiprom *et al*., 2024; Mitchum and Baum, 2008). The temporal precision of this silencing, coinciding exactly with cell wall dissolution and syncytium initiation, is consistent with effector-mediated transcriptional suppression, though the specific regulatory factors responsible remain to be identified. The progressive shift in WRKY, MYB, and bHLH transcription factor activity from strongly positive at 1 DPI to largely suppressed by 5 DPI mirrors the decline in phytoalexin accumulation and identifies these regulators as the most proximate candidates for follow-up functional characterization (Anguraj Vadivel *et al*., 2021; Bian *et al*., 2018; Liu *et al*., 2025; Wang *et al*., 2026).

The jasmonate pathway tells a parallel and equally informative story. Methyl dihydrojasmonate was constitutively suppressed from the moment of infection, not progressively declining but depleted at 45% of control levels at 1 DPI and remaining suppressed at all subsequent stages. A transient 12-hydroxyjasmonic acid elevation at 1 DPI (1.25× of controls) indicates a brief JA burst that is rapidly deactivated, and 13-oxoODE showed only a modest increase at 3 DPI without resulting in detectable JA-Ile accumulation. The JA pathway is not merely reduced but it is constitutively suppressed from the start of infection, consistent with active deactivation rather than passive downregulation and preceding syncytium formation (Erb and Reymond, 2019; Nahar *et al*., 2011; Zhang *et al*., 2017). The simultaneous constitutive suppression of JA and the sustained elevation of mevalonic acid lactone, which supplies isoprenoid precursors for cytokinin biosynthesis, creates the precise hormonal environment conducive to the nodulin induction and symbiosis program engagement observed at 5 DPI (Naseem and Dandekar, 2012). The simultaneous suppression of JA and elevation of cytokinin-relevant precursors therefore represents not two independent events but a coordinated hormonal reconfiguration that simultaneously dismantles defense and activates the symbiosis developmental program.

The temporal coincidence of trans-zeatin elevation at 1 and 5 DPI with nodulin-22 induction places our findings within the context of a decades-old observation now given its first integrated metabolomic-transcriptomic resolution. Cytokinin is a well-established inducer of nodulin gene expression during legume–rhizobium symbiosis (Dehio and de Bruijn, 1992; Lohar et al., 2004), and the elevation of trans-zeatin in SCN-infected roots at 1 and 5 DPI places the infected tissue in a hormonal state that is, by existing molecular precedent, conducive to nodulin gene activation. Whether SCN actively manipulates cytokinin levels through effector activity, or whether trans-zeatin elevation is an indirect consequence of syncytium-associated cell division and senescence suppression, the biological outcome is the same: infected tissue at 5 DPI exhibits the hormonal and transcriptional signatures of symbiosis-associated development. This convergence, cytokinin-relevant precursor elevation, nodulin induction, membrane remodeling lipids, and their co-regulation in the GC1-MC2 cluster, at the stage of active feeding site maturation provides the first integrated metabolomic-transcriptomic evidence for symbiosis program engagement in a compatible *Heterodera glycines*–soybean interaction.

The integrated metabolomic and transcriptomic analyses reported in this study demonstrate that SCN achieves compatibility by progressively dismantling host immunity across multiple coordinated pathways. Phenylpropanoid biosynthesis is suppressed downstream of PAL transcription, auxin homeostasis is disrupted through indole pathway redirection, jasmonate defense is attenuated from early infection, and by 5 DPI the legume symbiosis program is engaged through nodulin transcription and membrane lipid remodeling. This multi-pathway architecture explains the rapidity with which virulent SCN populations overcome *Rhg1*-mediated resistance (McCarville *et al*., 2017; Mitchum, 2016). The α-SNAP and NSF proteins encoded at *Rhg1* disrupt syncytium function in incompatible interactions (Bayless *et al*., 2016; Cook *et al*., 2012), but they represent a single molecular bottleneck. Our data shows that even in the presence of high-copy *Rhg1*, the virulent SCN successfully achieves compatibility through phenylpropanoid suppression, jasmonate attenuation, indole pathway redirection, and symbiosis program engagement, none of which require bypassing Rhg1. A single resistance locus cannot protect against a pathogen that dismantles immunity through coordinated multi-pathway suppression. Defense is nonetheless activated in compatible interactions, even in plants carrying high-copy *Rhg1*. The 27.9-fold corylin induction, the 3.1-fold coumesterol elevation, and the coordinated chalcone synthase and CYP450 induction at 1 DPI demonstrate that recognition occurs and defense begins. What fails is not initiation but its persistence through the syncytium initiation window, and this distinction matters for resistance engineering. Resistance strategies for amplifying early defense induction is therefore an insufficient strategy as the plant already mounts a robust initial response. The specific nodes at which compatible infection diverge from resistant interaction, including suppression of phenylpropanoid output downstream of PAL transcription, transcriptional collapse at 3 DPI, jasmonate attenuation, and symbiosis program engagement at 5 DPI, define the targets at which mechanistically grounded intervention would be most effective. Protecting phenylpropanoid output through the syncytium initiation window, maintaining jasmonate signaling through 3–5 DPI, and disrupting the nodulin induction associated with symbiosis program engagement represent engineering strategies that are grounded in mechanism rather than locus and go beyond what single-locus resistance can provide. The co-option of the legume symbiosis program by SCN suggests that legume vulnerability to sedentary endoparasitic nematodes may be an evolutionary consequence of the same developmental machinery that enables mutualistic endosymbionts (Geurts et al., 2016; Oldroyd, 2013). Understanding this convergence may therefore be essential for engineering crops that can distinguish friend from parasite at the cellular level.

## MATERIALS AND METHODS

### Plant growth and SCN inoculation

Soybean seeds of a high-copy *Rhg1* near-isogenic line LD09-15603 were surface-sterilized using vapor-phase ethanol sterilization (Kwon et al., 2024) and placed on moistened, autoclaved filter paper in 14-cm-diameter Petri dishes. Seeds were germinated in the dark at 22–24°C for 48 h. Two-day-old seedlings of uniform size were transplanted individually into steam-pasteurized sandy loam mix (2:1 v/v Torpedo sand: soil) in transparent polyvinyl carbonate tubes (17 cm × 2.5 cm diameter) set into plastic support trays (Miraeiz *et al*., 2026b). Plants were grown in a growth chamber at 27°C under a 16/8 h light/dark cycle and watered daily.

### Nematode culture and inoculation

A soybean cyst nematode (SCN) population of HG Type 2.5.7 was maintained on the susceptible cultivar Williams 82 at the Department of Crop Sciences, University of Illinois Urbana–Champaign. Eggs were harvested from stock plants and allowed to hatch following an established hatching protocol (Kwon *et al*., 2024). After seven days, pre-parasitic second-stage juveniles (J2s) were collected through a 25 µm sieve and adjusted to approximately 700 J2s/mL. The suspension was stirred gently on a magnetic stirrer for 2 minutes immediately before use to ensure uniform distribution. Three-day-old seedlings were inoculated by applying 1 mL of the J2 suspension directly to the root zone using a micropipette. Mock-inoculated control plants received 1 mL of sterile hatching solution without nematodes. Roots were harvested at 1-, 3-, and 5-days post-inoculation (DPI), immediately flash-frozen in liquid nitrogen, and stored at –80°C until further processing.

### RNA-extraction, library preparation and quality control

Total RNA was extracted from approximately 100 mg of finely powdered frozen root tissue using the E-Z® 96 Plant RNA Kit (Omega Bio-Tek) per the manufacturer’s instructions, followed by DNase I treatment (Thermo Scientific) to eliminate residual genomic DNA. RNA quality and integrity were verified prior to library preparation. mRNA-seq libraries were constructed using the Watchmaker mRNA-Seq Kit (Watchmaker Genomics) with polyA selection. Libraries were pooled and quantified by qPCR, then sequenced on a partial 25B lane of a NovaSeq X Plus instrument (V1.0 chemistry) at the Roy J. Carver Biotechnology Center, University of Illinois Urbana–Champaign. Paired-end sequencing was performed with 151 cycles per end, yielding 150 nt reads. The final dataset comprised three time points with four biological replicates per condition (SCN-inoculated and mock-inoculated). Raw read quality was assessed using FastQC (FastQC, 2015) (v0.12.1). Adapter trimming and quality filtering were carried out with fastp (Chen, 2025) (v1.0.1), with explicit Illumina TruSeq ligation adapter sequences provided for both reads and automatic adapter detection enabled for paired-end mode.

### Transcript quantification and differential expression analysis

Transcript-level abundance was quantified using Salmon (Patro et al., 2017) (v1.10.2) in selective-alignment mode against a decoy-aware index built from the Glycine max Wm82.a4.v1 reference transcriptome (Schmutz et al., 2010; Valliyodan et al., 2019). Quantification incorporated sequence composition and GC content bias corrections, with 100 bootstrap replicates per sample to support uncertainty-aware differential expression analysis. Transcript-level estimates were imported into R using tximport (Soneson et al., 2015) (v1.38.2) and summarized to gene-level counts.

A DESeq2 dataset was constructed with treatment as the sole factor in the design formula. Genes with fewer than 10 total counts across all samples were excluded. Differential expression analysis was conducted separately for each time point. DESeq2 (Love et al., 2014) (v1.50.2) estimated size factors from average transcript lengths supplied by tximport, fitted a negative binomial GLM, and applied Wald tests for pairwise contrasts between each treatment and the water control. Genes were considered differentially expressed at an adjusted p-value < 0.05 and |log₂ fold change| > 1 and were classified as either up– or down-regulated accordingly.

### Visualization and functional enrichment

For visualization purposes, a variance-stabilizing transformation (VST) was applied with dispersion estimation performed in a design-aware manner (blind = FALSE). Sample clustering was explored using t-distributed stochastic neighbor embedding (t-SNE) applied to the full VST-transformed gene set. Gene Ontology (GO) enrichment analysis for differentially expressed genes was performed using the GlycineMine (v5.1.0.3) web-based platform.

### WGCNA analysis

For the identification of modules in the transcriptome datasets, the 46,253 transcripts were used for WGCNA analysis in R. Hierarchical clustering based on the topological overlap (TO) similarity was used to obtain network modules, and the highly similar modules (dissimilarity < 0.25) were merged to prevent over-splitting of the modules. Module eigenvalues, representing the overall profile of the module, were calculated for each of the identified modules.

### Metabolite extraction and LC–MS analysis

Metabolites were extracted from ∼10 mg of freeze-dried, ground root tissue (pre-chilled in liquid nitrogen) using 25 µl/mg of ice-cold extraction solvent (methanol: water, 80:20 v/v) containing 0.1% formic acid and internal standards (2.5µM lidocaine). Samples were vortexed for 30 s, incubated overnight in a shaker at 4°C, sonicated briefly, and centrifuged at 16,000 g for 15 min at 4°C. The supernatant was collected, and the centrifugation step was repeated under the same conditions. The resulting supernatant was collected and stored at –80°C until LC–MS analysis.

LC–MS analysis was performed on a Waters Xevo G3 Q-ToF mass spectrometer coupled to an Acquity UPLC I-Class system (Waters Corporation, Milford, MA, USA). Chromatographic separation was achieved on an Acquity UPLC BEH C18 column (100 × 2.1 mm, 1.7 µm) maintained at 40°C. The mobile phase consisted of (A) water + 0.1% formic acid and (B) acetonitrile + 0.1% formic acid, with a 20-min gradient (5% B held for 1 min, increased to 95% B over 14 min, held for 2 min, then re-equilibrated for 3 min) at a flow rate of 0.3 mL min⁻¹. The Xevo G3 was operated in resolution mode with the following parameters: capillary voltage 3.0 kV (positive mode); sampling cone 40 V; source temperature 120°C; desolvation temperature 500°C; desolvation gas flow 800 L h⁻¹. Data were acquired in MS/MS mode: high collision energy ramp (15–50 V) for fragmentation. Mass accuracy was maintained using a continuous lock spray of leucine enkephalin ([M+H] + = 556.2771) at 200 pg µL⁻¹, infused at 10 µL min⁻¹.

Raw data files (.raw) were processed using MS-DIAL for peak detection, alignment, deconvolution, and quantification. Peak normalization was conducted by comparison with an internal standard (lidocaine). The acquired mass features were further used for Principal Component Analysis. Chemical identities were assigned to the mass-features by matching the MS^2^ fragmentation pattern with the MS^2^ ions of the corresponding metabolite reported in previous studies or as predicted by in-silico fragmentation by MS-DIAL, MS-FINDER, or SIRIUS following previously described method (Rai et al., 2020). Putative chemical identities were assigned to the acquired mass features by mapping onto the Kyoto Encyclopedia of Genes and Genomes (KEGG) pathway database with mass tolerance of 10 ppm.

### Multivariate analysis and DAM identification

Metabolite abundance data were log₂-transformed and Pareto-scaled to stabilize variance and reduce the influence of high-abundance features. Orthogonal partial least squares–discriminant analysis (OPLS-DA) was performed in SIMCA v17 (Sartorius) to maximize separation between SCN-infected and uninfected samples at each time point. Model validity was confirmed by cross-validation (Q² > 0.5) and permutation testing (n = 200 permutations; p < 0.05). Differentially abundant metabolites (DAMs) were identified by variable importance in projection (VIP) score > 1.5.

### Multi-omics integration

For integrative analysis, high-confidence annotated DAMs and DEGs across all time points were combined into a unified dataset. Pearson correlation coefficients were computed between normalized metabolite abundances and normalized gene expression values across all samples. Gene–metabolite pairs were considered strong if they met |r| ≥ 0.7, and *p* < 0.05. Hierarchical clustering (Ward’s linkage, Euclidean distance) was applied to identify co-regulated modules.

## DATA AVAILABILITY

The RNA-seq and metabolomics data supporting the findings of this study are available from the corresponding author upon request.

## AUTHOR CONTRIBUTIONS

M.R. and E.M. designed the study, performed the experiments, conducted the analysis, and wrote the original draft. M.D. helped with metabolome experiments. V.D. helped with sample preparation. M.H. and A.R. led the project, reviewed and edited the manuscript, and served as the corresponding authors.

## ACKNOWLEDGEMENTS

We extend our sincere gratitude to Alison Colgrove for providing SCN inoculum. We also thank Dr. Brian Diers for supplying the near-isogenic soybean lines used in this study.

## DECLARATION OF GENERATIVE AI AND AI-ASSISTED TECHNOLOGIES IN THE WRITING PROCESS

The authors used AI to improve the language of the manuscript and take responsibility for the content of the published article.

