## Supplementary figures and images for "Metabolic and transcriptional responses reveal co-option of symbiotic processes during early soybean-soybean cyst nematode interaction"

### Supplementary Figure S1

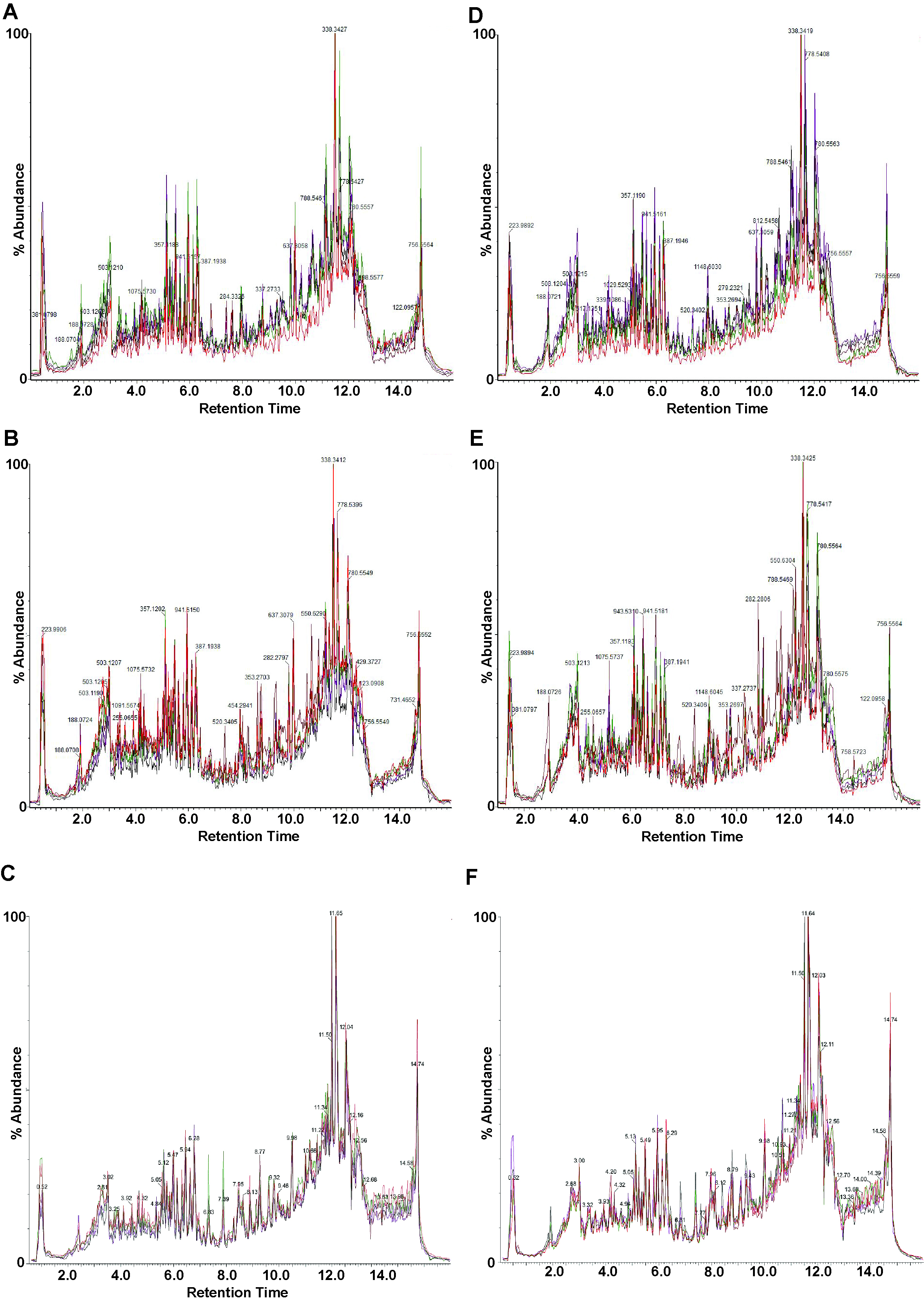

### Supplementary Figure S2

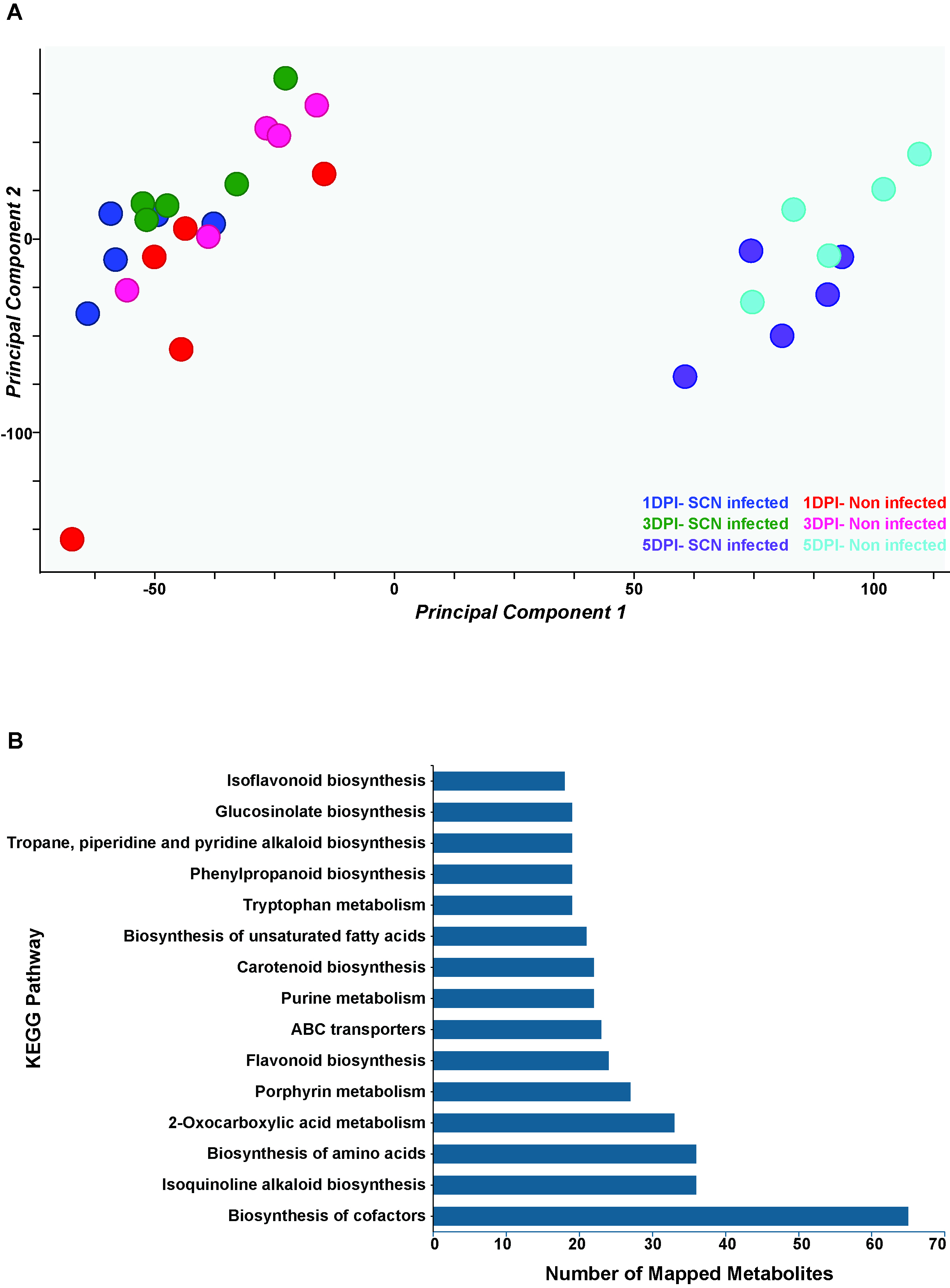

### Supplementary Figure S3

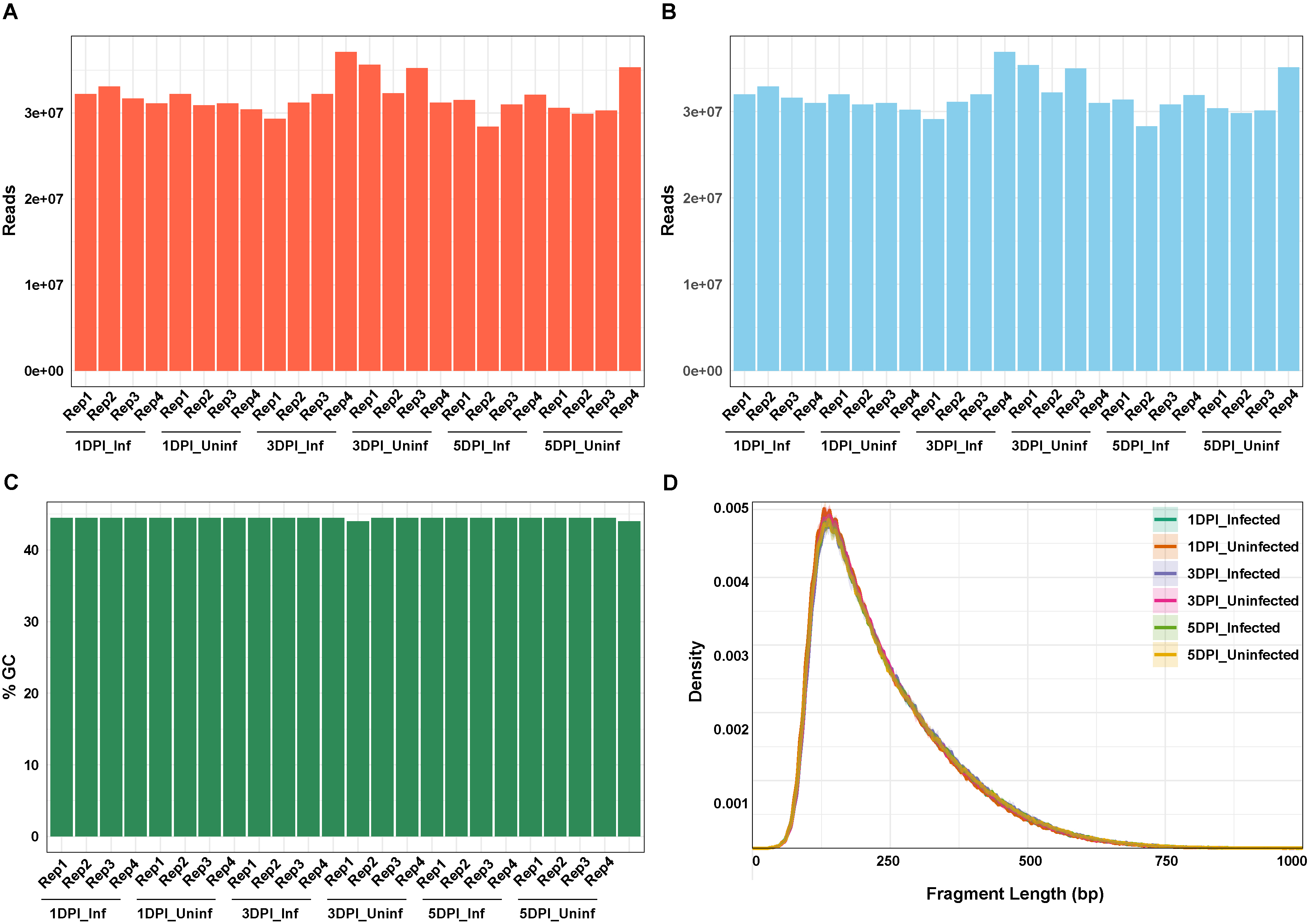

### Supplementary Figure S4

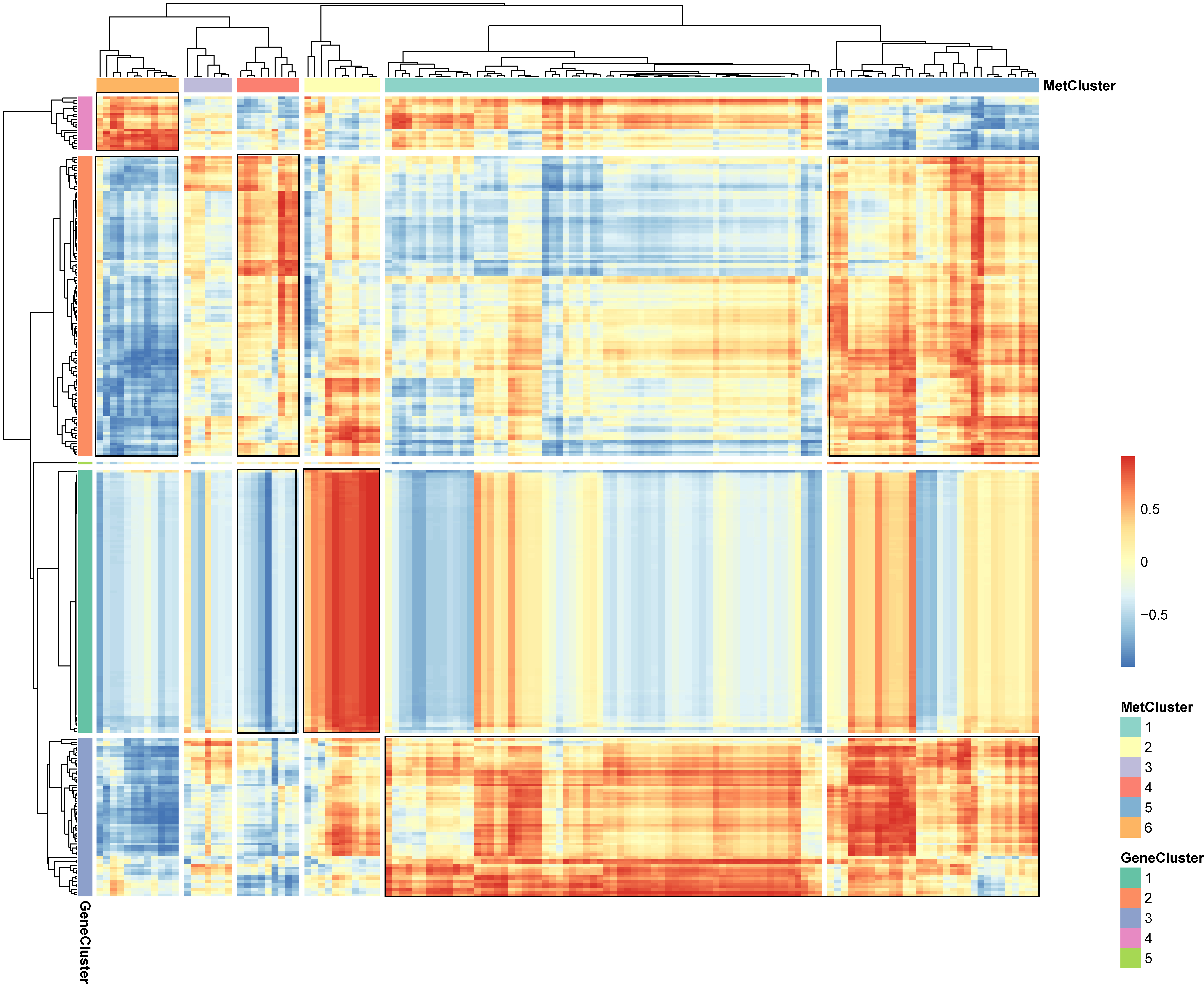

### Supplementary Figure S5

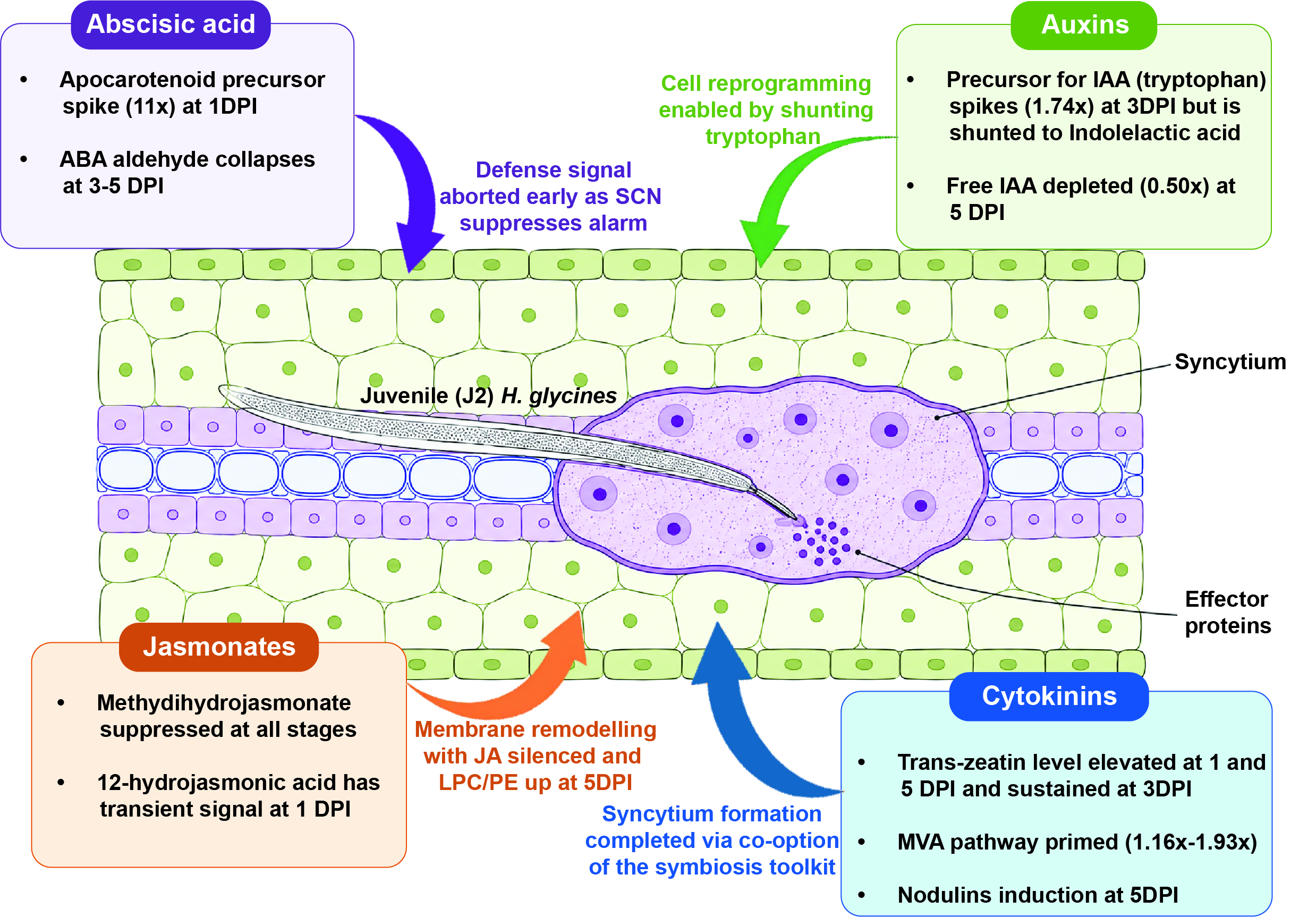

### Supplementary Figure S6

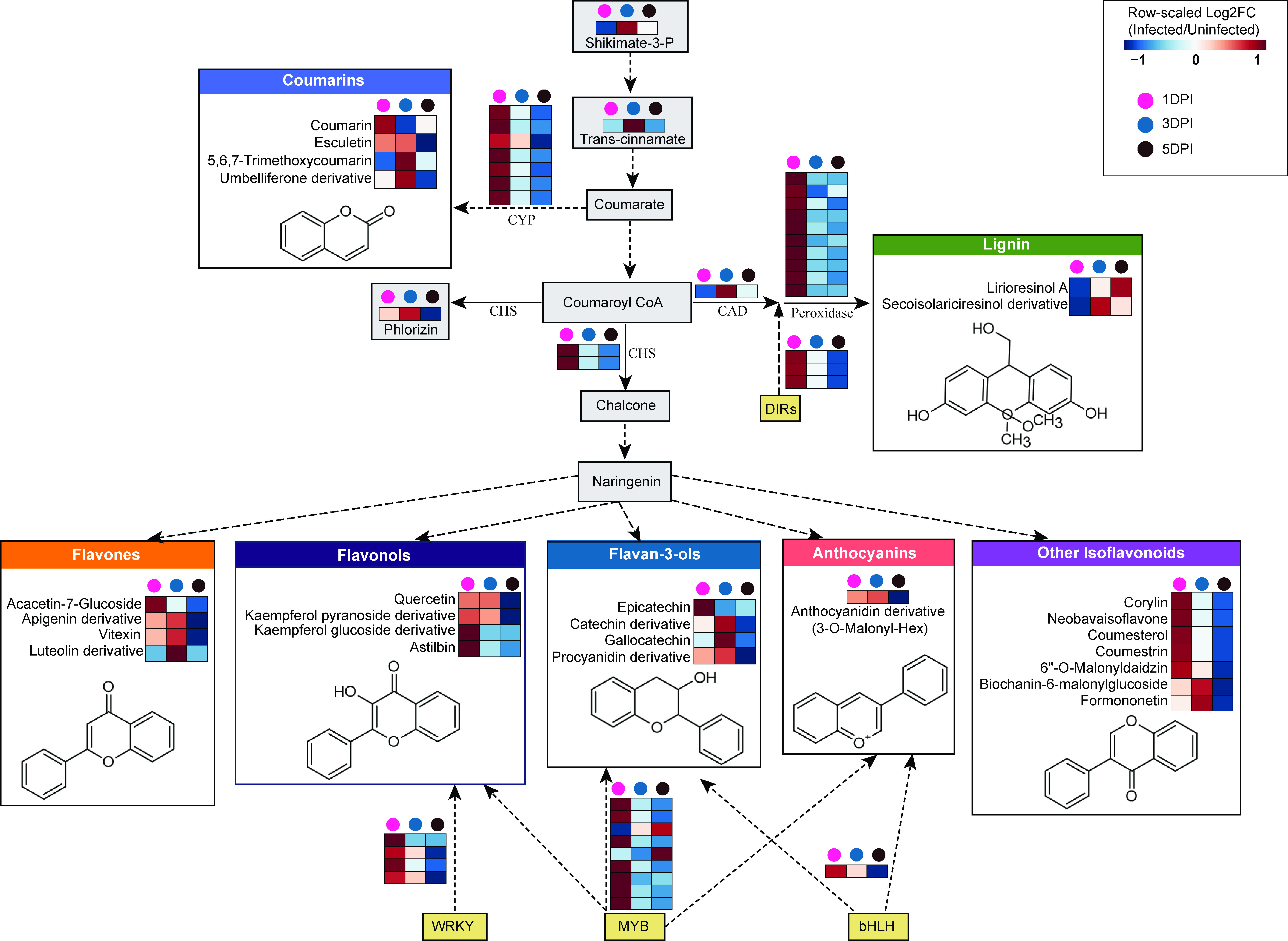
