## Supplemental Information for "Metabolic and transcriptional responses reveal co-option of symbiotic processes during early soybean-soybean cyst nematode interaction"

**^*^** Co-corresponding authors

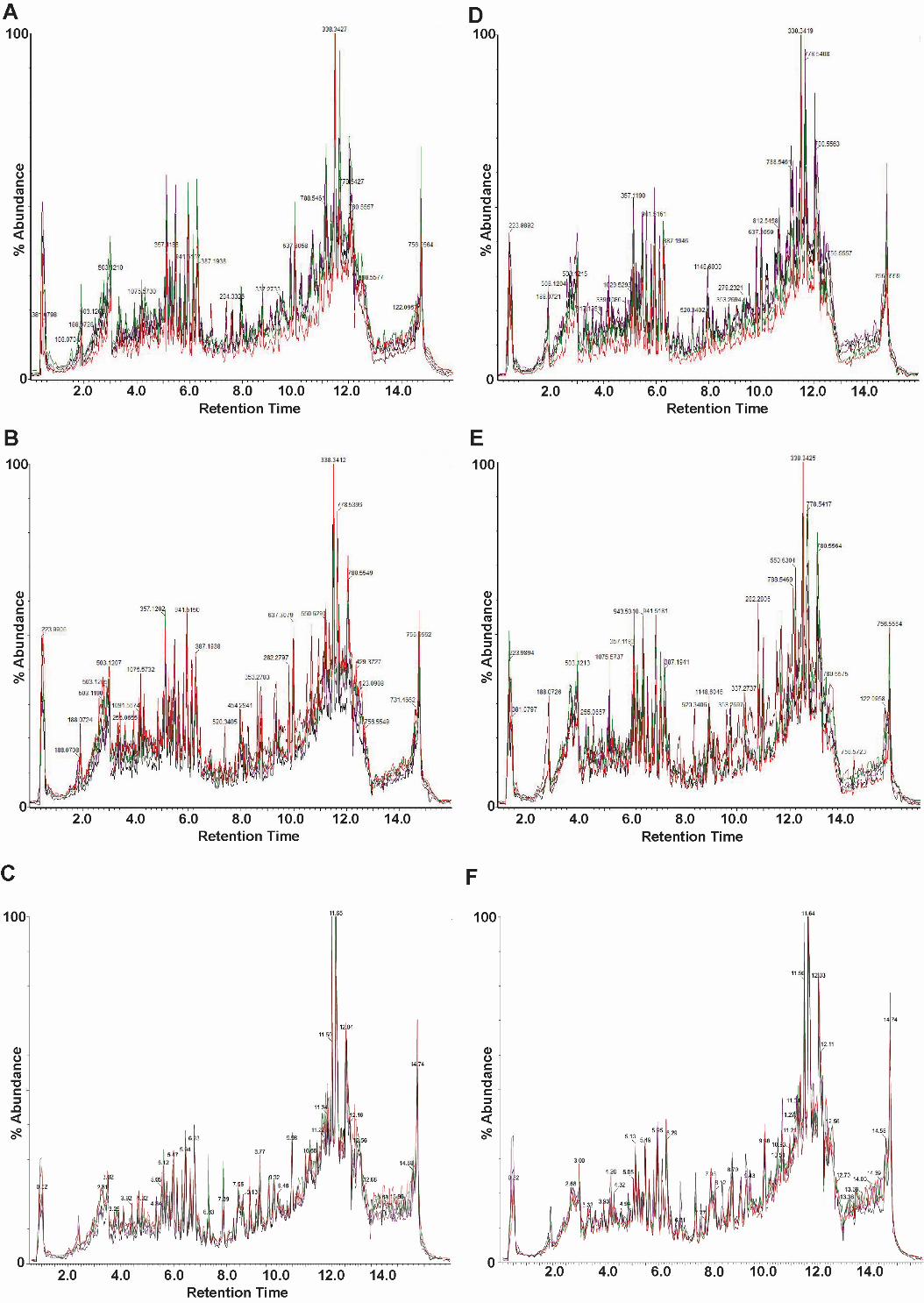

**Figure S1.** **Total ion chromatograms (TICs) of metabolomic profiles across treatments and time points.** Each panel represents one condition at a specific time point, with individual biological replicates overlaid.

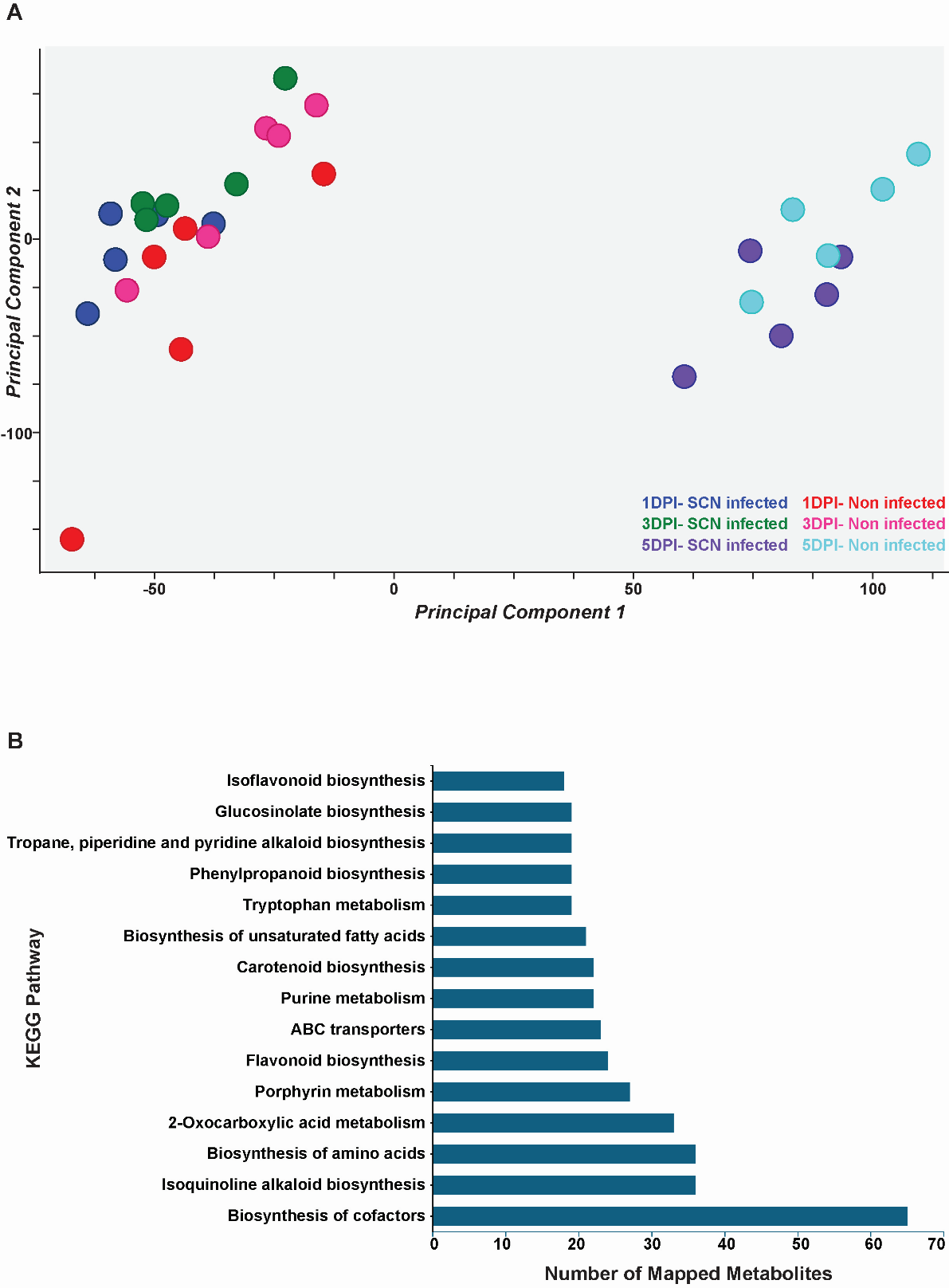

**Figure S2. Global metabolomic variation and KEGG-based pathway classification. (A)** Principal component analysis (PCA) of normalized LC–MS mass features. Each point represents a biological replicate. Percent variance explained by PC1 and PC2 is indicated on the axes. **(B)** Top 15 KEGG pathways ranked by the number of detected mass features assigned based on *m/z*. Bars indicate the total number of mapped features per pathway.

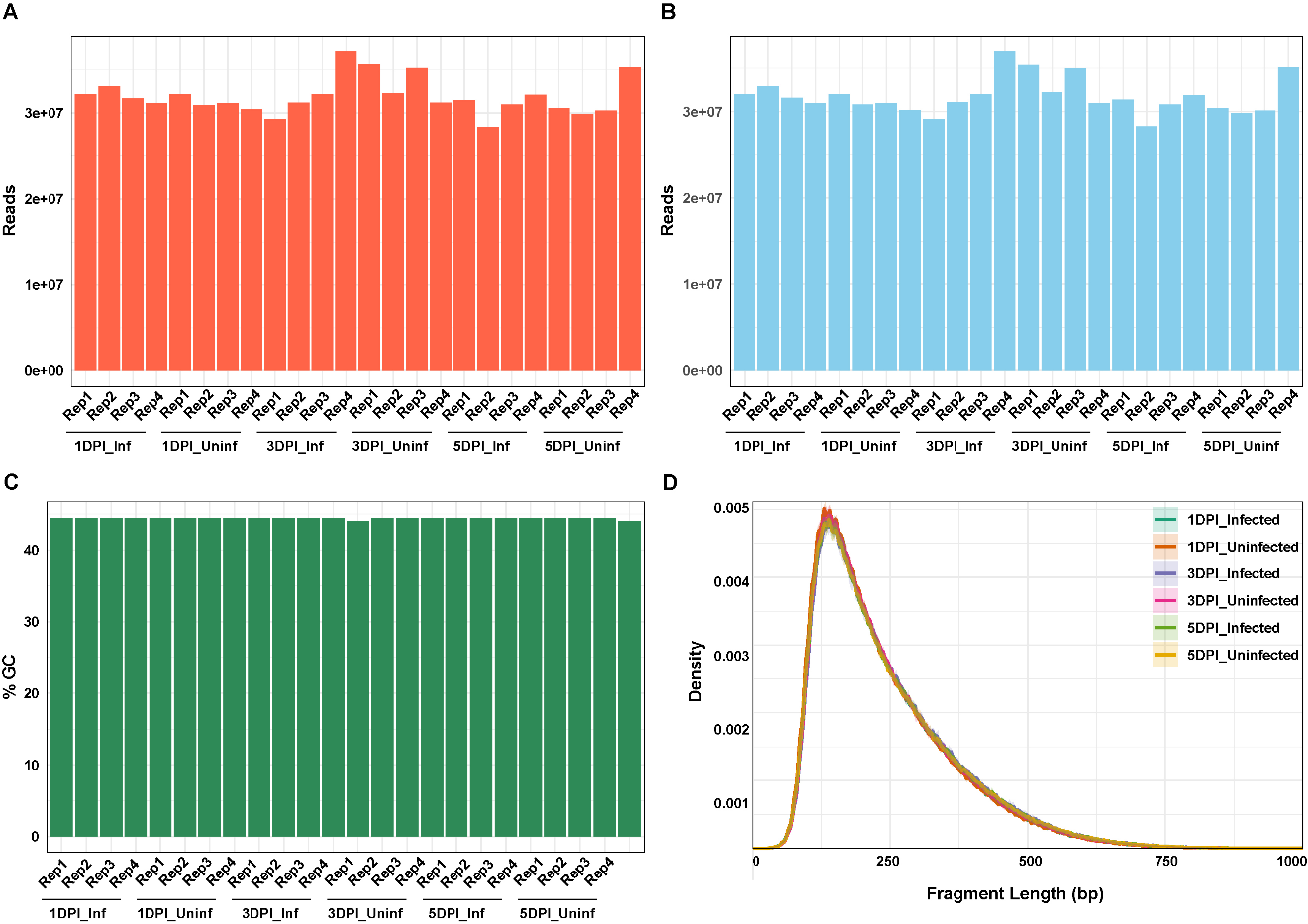

**Figure S3**. **Quality assessment of RNA-seq libraries across treatments and time points. (A)** Total number of raw reads generated per library for infected and non-infected samples at 1-, 3-, and 5-days post-inoculation (dpi). **(B)** Number of clean reads retained after quality filtering for each library. **(C)** GC content (%) of clean reads across all samples.
**(D)** Fragment length distribution of sequenced libraries showing consistent insert size profiles across treatments and time points.

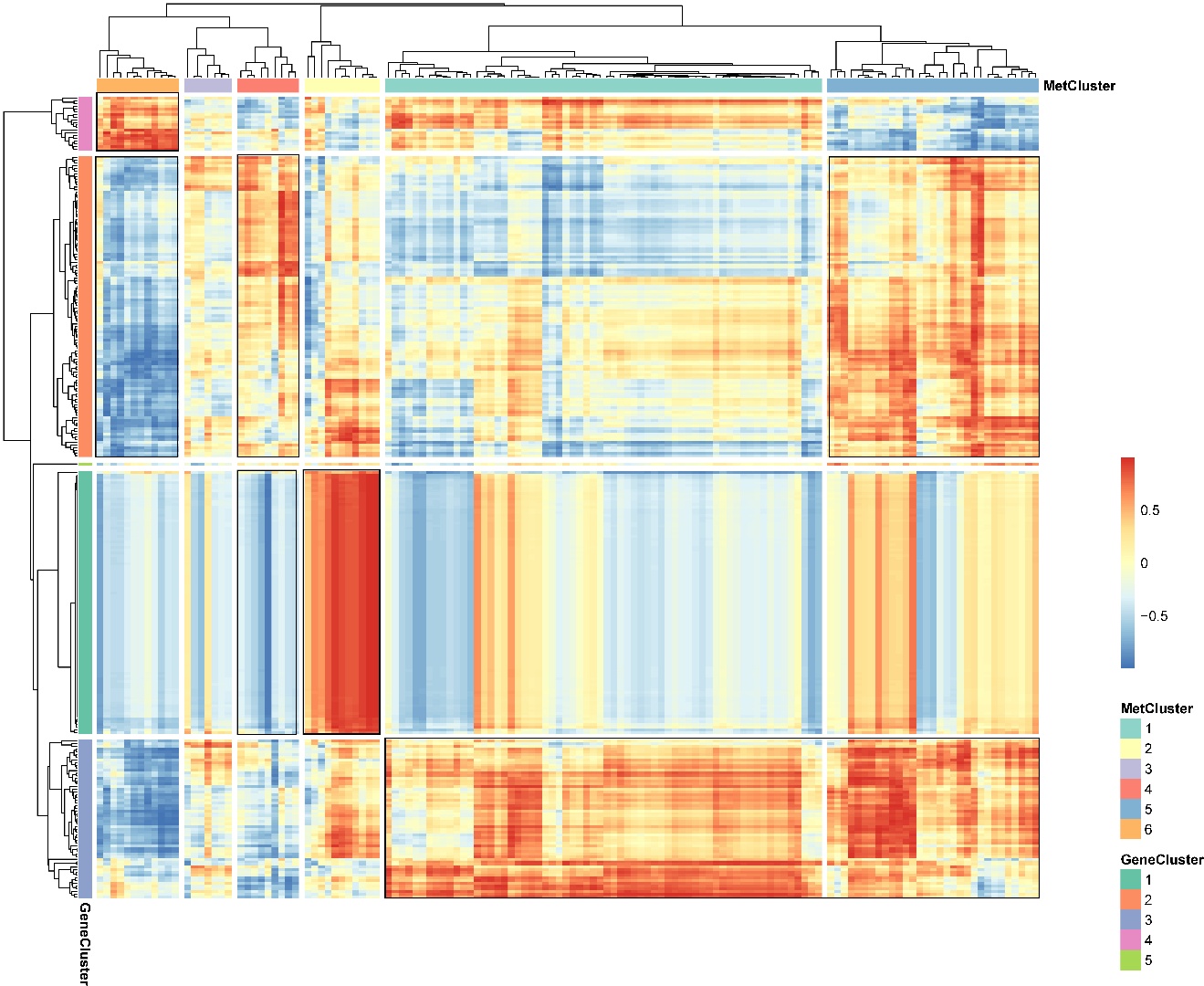

**Figure S4.** **Correlation analysis between differentially expressed genes (DEGs) and differentially accumulated metabolites (DAMs).** Heatmap showing pairwise correlation coefficients between DEGs and DAMs across all samples. Color intensity indicates the strength and direction of correlation (positive and negative).

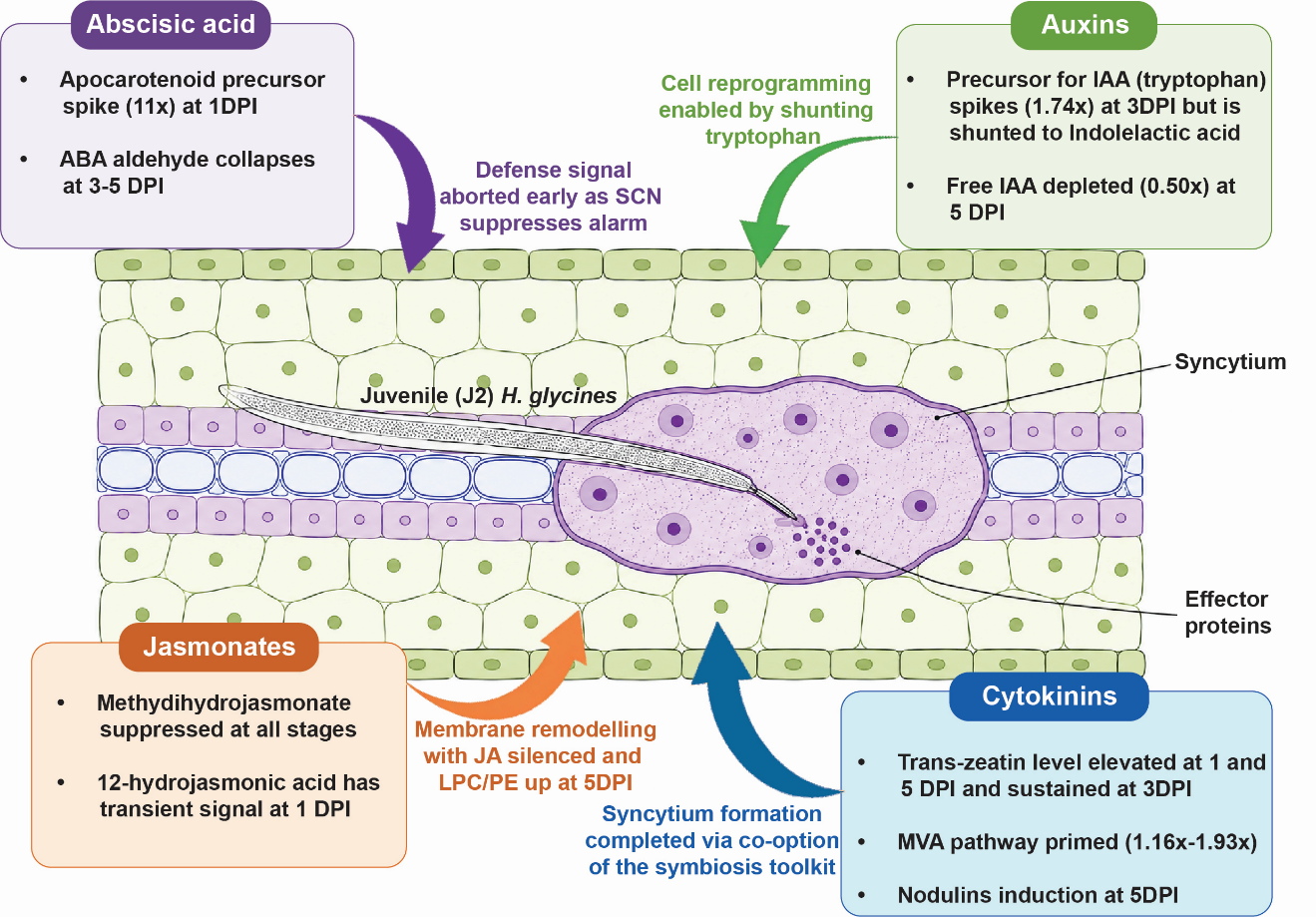

**Figure S5:** **Hormonal network reconfiguration during compatible soybean–*Heterodera glycines* interaction.** Stage-resolved hormonal transitions at 1-, 3-, and 5-days post-infection (DPI) in soybean roots carrying a high-copy *Rhg1* locus challenged with virulent SCN. The central root cross-section depicts the developing syncytium and attached *H. glycines* juvenile. ABA peaks transiently at 1 DPI before collapsing; free IAA is depleted by 5 DPI as tryptophan is shunted toward indolelactic acid; JA is suppressed across all stages; cytokinin is sustained at 3–5 DPI and associated with co-option of the symbiosis program at syncytium maturation.

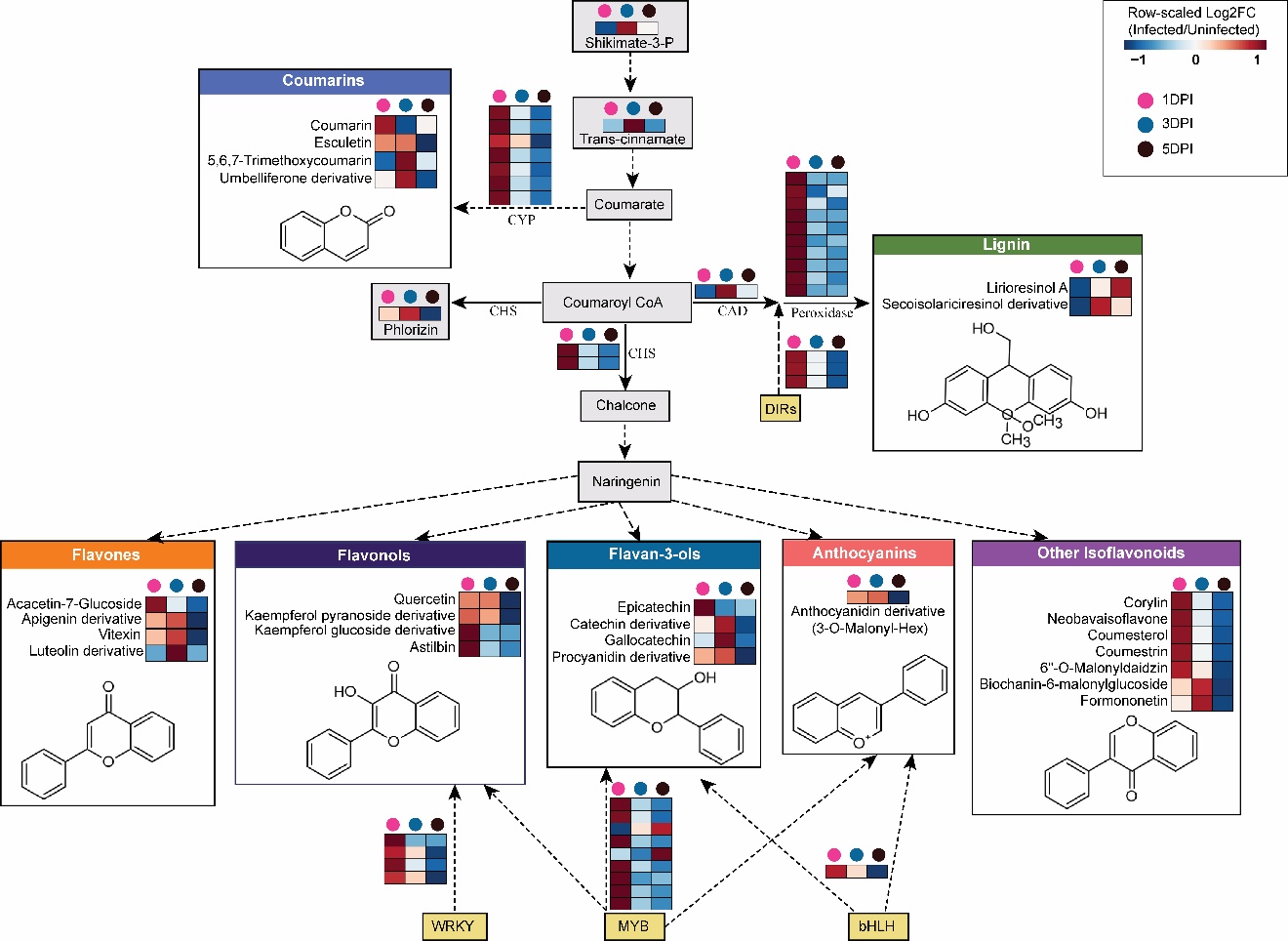

**Figure S6. Integrated transcript–metabolite pathway reconstruction based on highly correlated gene and metabolite clusters.** Genes and metabolites showing strong correlations between transcriptomic (gene clusters) and metabolomic (metabolite clusters) datasets were selected and mapped onto the phenylpropanoid and flavonoid biosynthesis pathways. Heatmaps represent row-scaled log2 fold change (infected/uninfected) at 1-, 3-, and 5-days post-inoculation (DPI). Colored circles above each heatmap indicate time points (pink, 1 dpi; blue, 3 dpi; dark gray, 5 DPI).
